# Invariant neural representation of parts of speech in the human brain

**DOI:** 10.1101/2024.01.15.575788

**Authors:** Pranav Misra, Yen-Cheng Shih, Hsiang-Yu Yu, Daniel Weisholtz, Joseph R Madsen, Sceillig Stone, Gabriel Kreiman

## Abstract

Elucidating the internal representation of language in the brain has major implications for cognitive science, brain disorders, and artificial intelligence. A pillar of linguistic studies is the notion that words have defined functions, often referred to as parts of speech. Here we recorded invasive neurophysiological responses from 1,801 electrodes in 20 patients with pharmacologically-resistant epilepsy while they were presented with two-word phrases consisting of an adjective and a noun. We observed neural signals that distinguished between these two parts of speech. The representation of parts of speech showed invariance across visual and auditory presentation modalities, robustness to word properties like length, order, frequency, and semantics, and even generalized across different languages. The selective signals were circumscribed within a small region in the left lateral orbitofrontal cortex. This selective, invariant, and localized representation of parts of speech extends classic fronto-temporal language models, providing a target for mechanistic and causal tests of how the brain represents basic building blocks of language, and introduces a systematic approach to elucidate the orchestration of more complex aspects of language.

## Introduction

Language plays a central role in almost all of our daily activities and is at the heart of how we interact with others^1,2^. Early neurological studies and subsequent work using electrical stimulation demonstrated that there exist specific brain regions that play essential roles in language understanding and production^3–8^. Despite the critical importance of language, progress towards elucidating the neural circuits underlying its representation has remained elusive, in part due to the difficulties in investigating animal models, and in part due to the challenges associated with examining neurophysiological responses in the human brain.

Several neurophysiological experiments have begun to investigate neural signals associated with presentation of individual words or short phrases^9–17^. There has been work examining the orthographic features of real versus pseudowords^18–20^, phonetic features of word comprehension^10,18,21,22^ and production^23–27^, retrieval of semantic information for audio-visual naming^10^, and semantic encoding^28^. These studies have shed light on the early processes associated with detecting, comprehending and producing words. Beyond individual words, at the heart of linguistic structures is the notion that words serve specific functions within a sentence, including articles, nouns, adjectives, and verbs. These parts of speech (POS) are widely shared across languages, are combined according to defined grammatical rules, and play critical roles in natural language processing algorithms^1,11,29–35^. Furthermore, recent work has suggested that POS may be implicitly learned and represented in modern large language models^36,37^. Many studies in patients with brain lesions have shown deficits in the retrieval of nouns versus verbs^17,19,38–42^. However, previous studies lacked the spatial and temporal resolution necessary to resolve explicit neural circuits associated with POS processing and did not evaluate the degree of invariance in the representation of POS.

What would a representation for parts of speech like nouns and adjectives in the brain look like? Consider the adjective “green” and the noun “apple”, combined to create the simple phrase “green apple.” Fundamental constraints for such a representation should include the basic invariances underlying the cognitive understanding of this phrase. The basic desiderata for the representation of parts of speech in language includes invariance to: (i) presentation modality (e.g., auditory versus visual), (ii) specific noun or adjective (e.g., green or red), (iii) position within a phrase (e.g., “green apple” versus “apple green”), (iv) specific language in bilingual speakers (e.g., “green apple” in English versus “manzana verde” in Spanish), (v) other word properties like their written length, number of syllables, and phoneme composition.

Here we set out to investigate the representation of parts of speech in the human brain by recording intracranial field potential responses with high spatiotemporal resolution and high signal-to-noise ratio from 1,801 electrodes implanted in 20 participants with pharmacologically-resistant epilepsy. We describe neural signals, especially in the left lateral orbitofrontal cortex, that selectively distinguish between nouns and adjectives. These selective signals for part-of-speech are robust when words are matched for orthography (e.g., word length), acoustic features (e.g., number of syllables), word sequence (e.g., noun or adjective at first or second position within a phrase), and frequency of occurrence. Remarkably, the representation of nouns versus adjectives generalizes across audio and visual modalities, across different semantic categories within each part of speech, and even across different languages.

## Results

We recorded intracranial field potentials from 1,801 electrodes (840 in gray matter, 961 in white matter) implanted in 20 participants with pharmacologically-resistant epilepsy via stereoelectroencephalography (**Table S1**). Participants heard (auditory modality) or read (visual modality) two words that were sequentially presented and indicated whether the words were the same or not (**Figure 1a**, **Methods**). Participants performed the task correctly on 93.6±7.7% (here and throughout unless stated otherwise, mean±SD) of the trials, which was significantly different from chance level (p<0.01, two tailed t-test). All electrode locations are shown in **Figure 1b-g** (see also **Tables S1-S2** and **Methods**). We used a bipolar reference, and we focus on the intracranial field potential signals filtered in the high gamma frequency band, referred to as neural responses throughout and reported in the plots as normalized gamma power (65-150 Hz, **Methods**). Audio duration and word length for all stimuli are shown in **Figure 1h–j**.

**Fig. 1.**
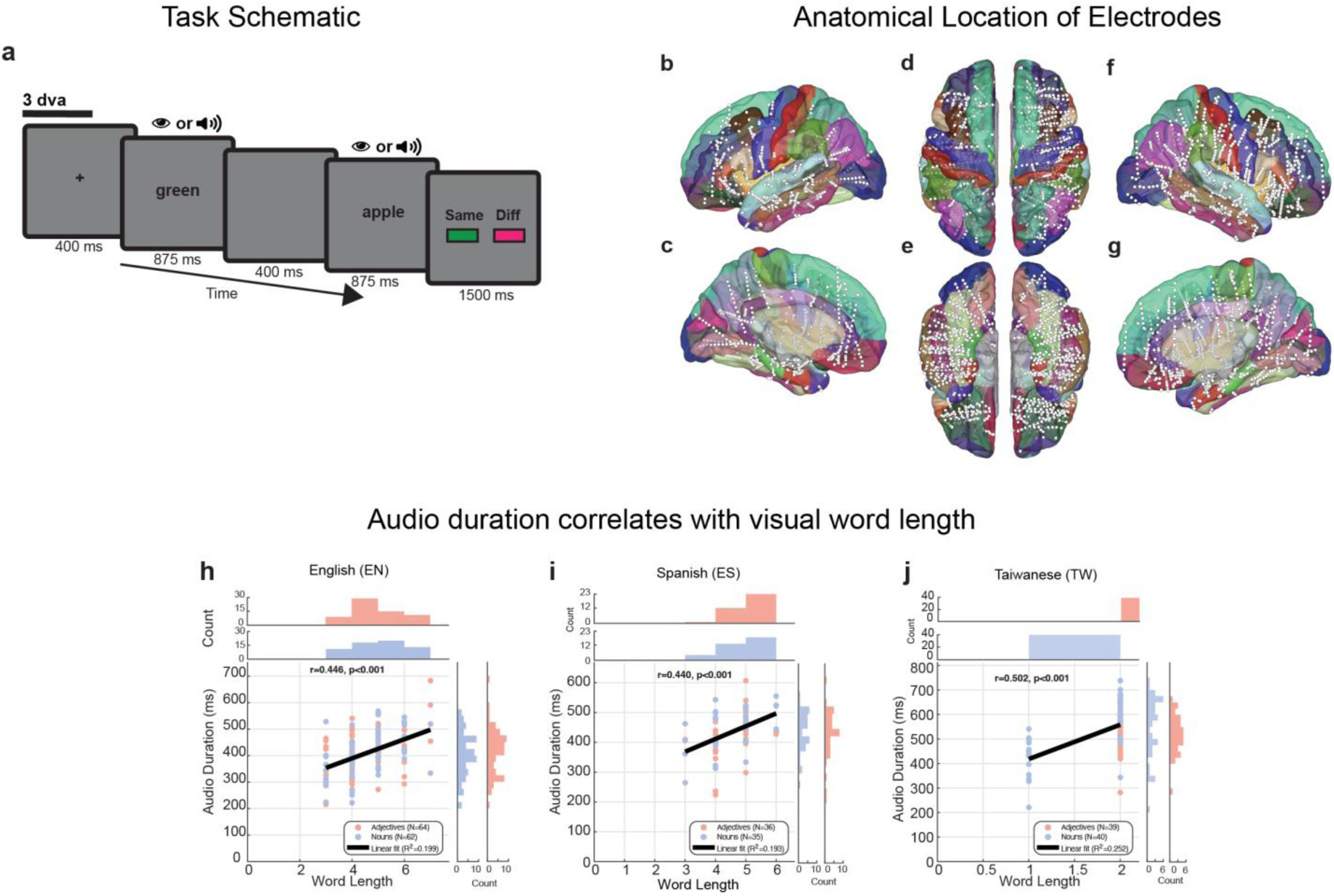
Task design and electrode locations. **a.** Two words were sequentially presented either in visual modality or auditory modality. Participants indicated whether the two words were the same (e.g., “apple apple” or “green green”, 8% of trials of each type) or different (e.g., “green apple” or “apple green”: 42% of trials of each type, Methods). In the 84% of trials where the two words were different, there was an adjective followed by a noun or a noun followed by an adjective. **b-g**. Location of all electrodes overlayed on the Desikan-Killiany Atlas shown with different views. Each white circle shows one electrode. **b**. Left lateral view (n=693), **c**. Left medial view (n=693), **d**. Superior, whole brain view (n=1,801), **e**. Inferior, whole brain view (n=1,801), **f**. Right lateral view (n=1,108) **g**. Right medial view (n=1,108). **h-j.** Scatter of audio word duration (y-axis) in milliseconds and word length (x-axis) for nouns (blue) and adjectives (red) for English **(h),** Spanish **(i),** and Taiwanese words (**j**). For English, Spanish, and Taiwanese, mean audio durations were 421 ± 86 ms, 438 ± 72 ms, and 538 ± 103 ms, respectively. Nouns averaged 426 ± 89 ms (EN), 443 ± 70 ms (ES), and 549 ± 124 ms (TW), while adjectives averaged 414 ± 81 ms (EN), 434 ± 80 ms (ES), and 527 ± 81 ms (TW). Mean text lengths were 4.8 ± 1.2 characters (EN), 4.6 ± 0.8 characters (ES), and 1.8 ± 0.4 characters (TW). For nouns, text lengths averaged 4.9 ± 1.2 characters (EN), 4.5 ± 0.8 characters (ES), and 1.7 ± 0.5 characters (TW); for adjectives, text lengths averaged 4.5 ± 1.1 characters (EN), 4.7 ± 0.7 characters (ES), and 2.0 ± 0.0 characters (TW). Duration correlated with text length in all three languages (R² = 0.19–0.25, p < 0.001). Word-type comparisons between nouns and adjectives showed no differences in audio duration for any language (p>0.05, t-test) (**Methods**).

### Neural signals reflect visual, auditory and multimodal inputs

We observed 565 electrodes (31.4% of the total) that responded to auditory stimuli (**Figure S1a-c, g-i**, **Methods**) and 532 electrodes (29.5% of the total) that responded to visual stimuli (**Figure S1d-f, g-i**). The overall proportions and dynamics of visual and auditory responsive signals are consistent with previous work^24,43^. Of these electrodes, there were 293 electrodes that responded to *both* auditory and visual stimuli (**Figure S1g-i**). These 293 electrodes represent 16.3% of the total, 51.9% of the auditory responsive electrodes, and 55.0% of the visually responsive electrodes. This number of audiovisual electrodes is unlikely to arise by chance from

the number of auditory and visual electrodes (p<10^-4^, permutation test, n=10^6^ iterations). Of these 293 electrodes, 147 (50.2%) were in the left hemisphere and 146 (49.8%) were in the right hemisphere. **Figure S1j-m** shows the responses of an example audiovisual responsive electrode located in the left rostral middle-frontal gyrus (**Figure S1m**). This electrode showed strong evoked responses evident in the trial-averaged responses (**Figure S1j**), and even in individual trials for both auditory stimuli (**Figure S1k**) and visual stimuli (**Figure S1l**).

To compare the response dynamics of auditory and visual responses, we calculated the time at which the neural signals reached half of the max amplitude (half-maximum time, arrows in **Figure S1j**, **Methods**) for neural responses such as those in **Figure S1j**. There was no significant difference between the half-maximum time for auditory-only electrodes (329±187 ms, grey-left in **Figure S1n**) and visual only electrodes (336±174 ms, black-middle in **Figure S1n**) (p>0.05, ranksum test). Similarly, there was no significant difference between the half-maximum time for the audio and visual responses of audiovisual electrodes (379±193 ms versus 341±174 ms, half-black-right in **Figure S1n**, p>0.05, ranksum test). However, there was a small but significant difference between the half-maximum time for audio only electrodes and auditory responses of audiovisual electrodes (p<0.01, ranksum test). As expected, for the audio-only electrodes, the response area under the curve (AUC) for auditory stimuli (108±100 µV^2^/Hz-ms) was larger than for visual stimuli (44±16 µV^2^/Hz-ms) (p<10^-4^, ranksum test, **Figure S1o**). Similarly, for the visual-only electrodes, the response AUC to auditory stimuli (40±23 µV^2^/Hz-ms) was smaller than to visual stimuli (53±43 µV^2^/Hz-ms) (p<10^-4^, ranksum test, **Figure S1p**). For the audiovisual electrodes, the response AUC to auditory stimuli (71±72 µV^2^/Hz-ms) was slightly larger than to visual stimuli (54±39 µV^2^/Hz-ms) (p<0.01, ranksum test, **Figure S1q**).

We compared the response effect sizes for visual versus auditory responses for all electrodes (**Figure S2a**, **Methods,** example electrode from **Figure S1j-m** shown with a green arrow). As expected from the electrode definitions, for the audio-only electrodes (cyan hexagrams), the effect size for auditory stimuli (19.8±5.4%) was larger than for visual stimuli (10.0±4.4%, p<0.01, ranksum test). Similarly, for the visual-only electrodes (black squares), the effect size for visual stimuli (19.0±4.8%) was larger than for auditory stimuli (10.2±4.3%, p<0.01, ranksum test). For the audiovisual electrodes (orange triangles), the effect size for auditory stimuli (19.3±5.5%) was not different from the one for visual stimuli (19.2±4.8%, p>0.01, ranksum test). A similar scatter plot for the rostal middle frontal region is shown in **Figure S2b**. Of the 41 regions in the Desikan-Killiany Atlas where we had sampling (34 defined regions and 7 extra regions representing deep gray matter structures, **Methods**, **Figure 1**, **Tables S1-S2**), 13 regions had a significantly higher number of multimodal electrodes than expected by chance from the number of audio and visual electrodes (p<0.01, permutation test, n=10^6^ iterations, **Figure S2 a-c,** indicated in bold in **Table S2**).

In sum, we observed electrodes that responded exclusively to auditory stimuli, other electrodes that focused exclusively on visual stimuli, and also other electrodes that revealed multimodal responses to both auditory and visual inputs. On average, there were large amplitude differences for unimodal electrodes but only subtle differences in the latencies and amplitudes of responses to auditory versus visual stimuli for the multimodal electrodes. Responsiveness here is characterized descriptively; subsequent analyses of selectivity use independent criteria.

### Multimodal neural signals distinguish different parts of speech

We evaluated whether the neural signals differentiated between nouns and adjectives. Nouns and adjectives were matched for their number of syllables and word length to control for potential confounds not specific to parts of speech (**Table S3**, **Methods**). **Figure 2** shows the responses of an example electrode located in the orbital H-shaped sulcus within the left lateral orbitofrontal cortex (**Figure 2i** depicts the electrode location). The orbital H-shaped sulcus lies above the bone of the eye socket where a butterfly-like gyrus can be seen, formed along H-shaped recessions of the sulcus. The neural responses are aligned to the word onset (vertical dashed line) for auditory presentation (**Figure 2a, b**) or visual presentation (**Figure 2c, d**), for the first (**Figure 2a, c**), or second (**Figure 2b, d**) word in each trial. This electrode showed multimodal responses triggered by both auditory and visual stimuli. The responses to nouns (blue) were stronger than adjectives (red) across all four conditions, including both word 1 and word 2, and both for visual and auditory stimuli. The differences between nouns and adjectives can be readily appreciated even in individual trials (**Figures 2e-h**). These differences became significant at approximately 430 ms after word onset for visual presentation and about 610 ms for auditory presentation.

**Fig. 2.**
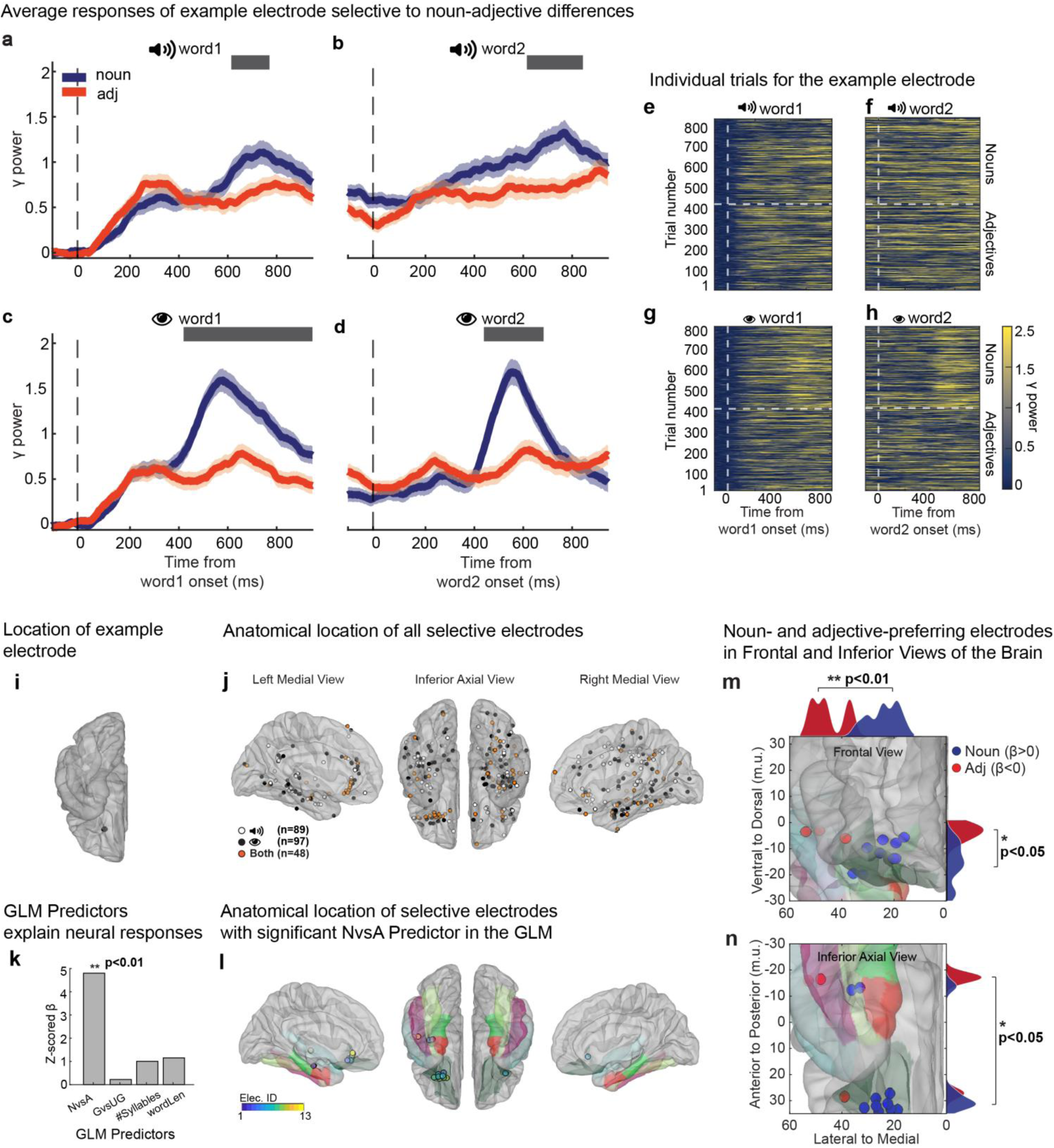
Neural signals distinguish between different parts of speech. **a-d.** Trial-averaged normalized gamma-band power of responses from an example electrode in the left lateral orbitofrontal cortex (see location in **i**) to nouns (blue) or adjectives (red) during presentation of auditory stimuli (**a, b,** n=435 grammatical and 432 ungrammatical trials) or visual stimuli (**c**, **d,** n=435 grammatical and 432 ungrammatical trials) aligned to the onset (vertical dashed line) of the first word (**a**, **c**) or second word (**b**, **d**). Shaded areas denote s.e.m. Horizontal gray lines denote windows of statistically significant differences between responses to nouns versus adjectives (t-test p<0.05, Benjamini-Hochberg false detection rate, q<0.05). **e-h**. Raster plots showing the responses in each individual trial (see color scale on bottom right). The blue and red curves in **a-d** correspond to the averages of noun and adjective trials, respectively, in **e-h. i** Location of the example electrode in the left lateral orbitofrontal cortex. **J** Electrodes that were selective to differences between nouns and adjectives for audio-only (white; n=89; left=30; right=59), visual-only (black; n=97; left=41; right=56) and multimodal (orange; n=48; left=21; right=27) stimuli. Brain is shown in three views: left medial (left, n=92), inferior axial (center), and right medial views (right, n=142). **k.** Z-scored β coefficients for Generalized Linear Model used to predict area under the curve between 200 ms and 800 ms post word onset, using four task predictors: Noun versus Adjectives, Grammatically correct versus incorrect, number of syllables (auditory presentation) and word length (visual presentation). Asterisks denote statistically significant coefficients, corrected for multiple comparisons (**Methods**). **l.** Electrodes that revealed statistically significant differences between nouns and adjectives for both audio and visual presentation (n=13 electrodes; colors indicate unique electrodes to match them across the different brain views). Electrodes whose responses were significantly explained only by the Nouns versus Adjective task predictor in the GLM are included in this plot. iELVis pullout factor=0, opaqueness=0.4. The fusiform (light green), parahippocampal (lime yellow), LOF (dark green-gray), entorhinal (red), inferior-temporal (magenta) and superior-temporal (cyan) regions are shown in the color scheme of the Desikan-Killiany Atlas (Figure 1b-g, Methods). **m,n.** All electrodes from **l** projected onto the left hemisphere are shown on the frontal plane (**m**) and the axial plane (**n,** same plane as **l**). All the electrodes that respond more strongly to nouns, i.e., Nouns versus Adjectives β>0 (n=10 electrodes), are shown in blue and electrodes that responded more strongly to adjectives (β<0, n=3 electrodes), are shown in red. All units are in MNI305 coordinates. Kernel density curves (bandwidth 2) outline the marginal distributions of noun-preferring (blue) and adjective-preferring (red) electrodes along the medial-lateral axis (**m,n:** x-axis, zero being more medial), ventral-dorsal axis (**m:** right z-axis) and posterior-anterior axis (**n:** right y-axis). P-values indicate significant differences between the coordinates for noun- and adjective-preferring electrodes (ranksum test).

In all, there were 89 electrodes (**Figure 2j**, white), 97 electrodes (**Figure 2j**, black), and 48 electrodes (**Figure 2j**, orange; total=234) that showed a difference between nouns and adjectives for auditory stimuli only, visual stimuli only, or both modalities, respectively (p<0.05, t-test, Benjamini-Hochberg false detection rate, q<0.05, calculated per modality, **Methods**). These electrodes represent 24% (89+48=137 out of 565), 27% (97+48=145 out of 532), and 16% (48 out of 293), of the auditory responsive, visually responsive, and multimodal responsive electrodes, respectively. It is unlikely that the 48 multimodal electrodes distinguishing nouns and adjectives could be ascribed to randomly sampling from the total of auditory and visual electrodes (p<10^-4^, permutation test, n=10^6^ iterations). **Figure S2h** shows the regions that had a larger number of multimodal electrodes distinguishing nouns versus adjectives compared to the null hypothesis obtained by randomly sampling from the total of auditory and visual electrodes, like the lateral orbitofrontal cortex shown in **Figure S2e,g** (p<10^-4^, permutation test, n=10^6^ iterations, see **Figure S3a** for a schematic of the sequential evaluation steps and electrode count at each step).

Next, we characterized the dynamics and effect size for selective responses. We calculated the selectivity response window (max window) as the length of the longest continuous period with a significant difference between nouns and adjectives across both words (**Figure S2d,** similar to gray horizontal bars in **Figure 2a-d**) and the effect size as the percentage change in the response with respect to baseline (**Figure S2f**). As expected, the max window and effect sizes for unimodal electrodes were consistent with their modality preferences (**Figure S2d, f**; cyan: audio, black: vision) and were enhanced for both vision and audio in the multimodal electrodes (**Figure S2d, f**; orange). The electrodes in the lateral orbitofrontal cortex are separately shown in **Figure S2e** and **S2g** (the example electrode from **Figure 2a-h** is shown with a green arrow).

In sum, we observed electrodes that selectively distinguished between nouns and adjectives, both for auditory stimuli and visual stimuli, and irrespective of the specific order in which the noun and adjective were presented.

### Neural selectivity for nouns versus adjectives was robust to word properties, phrase grammar, usage frequency, and word subcategory

Even though nouns and adjectives were matched in their average number of syllables and word length, we asked whether these variables could still contribute to the neural responses differentiating nouns and adjectives. Additionally, each trial could be grammatically correct (e.g., “green apple”), or incorrect (e.g., “apple green”) (**Methods**). Therefore, we asked whether grammar could contribute to the neural differences between nouns and adjectives. To address these questions, we built a generalized linear model (GLM) for each electrode to predict its response AUC between 200 ms and 800 ms after word onset using four predictors: nouns versus adjectives, grammatically correct or not, word length (vision) or number of syllables (audition) (**Methods**). The predictor coefficients in the GLM model for the example electrode in **Figure 2a-d** show that only the nouns versus adjectives label significantly explained the neural responses for both auditory and visual presentation (**Figure 2k**). A total of 14 electrodes showed nouns versus adjectives as the *only* statistically significant predictor in the GLM analysis; 13/14 (93%) of these electrodes distinguished nouns versus adjectives for both auditory and visual inputs, such as the example electrode in **Figure 2a-i** (one additional POS visual-only, **Figure S4k-t**).

**##Table S4** shows the distribution of electrodes distinguishing part of speech as the only GLM predictor between the left and right hemispheres for all brain regions and **Table S5** shows the distribution of electrodes separating nouns versus adjectives as the only GLM predictor in different participants. The locations of electrodes that robustly distinguished nouns and adjectives as the only predictor in the GLM (**Figure 2l**) revealed a cluster enriched in the left lateral orbitofrontal cortex (LOF). Within the left LOF, all of the electrodes (8 out of the 8) were in the posterior part of the orbital H-shaped sulcus. We recorded from a total of 113 electrodes in the lateral orbitofrontal region, 38 electrodes in the left hemisphere and 75 electrodes in the right hemisphere (**Figure 1b-g, Table S2**). **Figure S3b** shows the number of electrodes along each step of processing for the lateral orbitofrontal region (see **Figure S3a** for the corresponding number for all regions combined). Of the 38 left hemisphere LOF electrodes, 21% distinguished nouns from adjectives during both audio and visual presentation. In stark contrast, only 1.3% of the 75 LOF electrodes in the right hemisphere distinguished nouns from adjectives in both audio and vision (these hemispheric differences were statistically significant: p<10^-4^, permutation test, n=10^6^ iterations).

We had hypothesized that distinguishing parts of speech constitutes a core component of language and would therefore be reflected *exclusively in both* visual and auditory modalities. Indeed, 13/14 (93%) of electrodes differentiating nouns from adjectives in the GLM did so in both modalities. In addition to these 13 electrodes there was a small number of electrodes (2 auditory only and 1 visual only) that showed differences between nouns and adjectives in one modality but not the other. Unlike the electrodes in **Figure 2l**, for the 2 auditory-only electrodes, the number of syllables also significantly contributed towards explaining the neural responses. **Figure S4a-j** shows the responses of an example electrode located in the right insula that showed a difference between nouns and adjectives during auditory presentation but *not* during visual presentation. The number of syllables also contributed to the response prediction for this electrode (**Figure S4j**). Conversely, **Figure S4 k-t** shows the responses of an example electrode located in the left lateral orbitofrontal cortex, but outside the posterior part of the orbital H-shaped sulcus, that showed a clear difference between nouns and adjectives during visual presentation but *not* during auditory presentation. This electrode showed a small difference between nouns and adjectives also in the auditory modality, but this difference was not statistically significant. **Figure S4 u,v** shows the locations of auditory only (white circles) and visual only electrodes (black circle) in the left (**u**) and the right hemispheres(**v**) distinguishing between nouns and adjectives.

In all, the responses of 52 out of 234 electrodes were predicted by at least one variable in the GLM (**Figure S3a**). For 16 out of 52 electrodes (31%), parts of speech was a significant predictor, the responses of the other 36 electrodes were correlated with other variables. Of those 16 electrodes, the responses of 14 electrodes (88%) were explained by parts of speech only (**Table S4)**, and for the remaining 2 the number of syllables was also a significant predictor (**Figure S4u,v,** see also **Figure S3 a,b**). **Figure S3c** shows which predictors in the GLM significantly explained the non-part-of-speech responses in **Figure S3a.**

We considered the possibility that using separate GLMs for each modality may reveal electrodes that better capture the contributions from other predictors. **Figure S5a,b** shows the modality specific GLMs for the example electrode in **Figure 2a-d**, separately for auditory (**Figure S5a**) and visual stimuli (**Figure S5b**). Part of speech was the only GLM predictor that significantly explained the neural responses of this electrode for both modalities. With this dual approach, there was a slight increase in the number of electrodes that were modulated by other predictors in addition to parts of speech (**Figure S5c**; electrode counts per predictor combination, pooled across modality-specific GLMs). However, the number of electrodes whose responses were accounted for by parts of speech only remained similar to the one with the combined GLM analyses (n=13, **Figure S5c;** see electrode locations in **Figure S5d**).

Nouns and adjectives differ in their usage frequency. We asked whether the differences in the neural responses to nouns versus adjectives depended on usage frequency. To address this question, we randomly subsampled the trials to match the distribution of Google Books Ngram frequency (**Methods**). The Google Books Ngram documents the usage frequencies of words in printed sources published between 1500 and 2022. **Figure S6a** shows matched noun and adjective distributions for the example electrode shown in **Figure 2a-i,k.** This electrode showed differential responses between parts of speech for auditory (**Figure S6b,c,f,g**) and visual (**Figure S6d,e,h,i**) stimuli during word1 (**Figure S6b,d,f,h**) and word2 (**Figure S6c,e,g,i**), even after nouns and adjectives were matched for their frequency of occurrence. Of the 13 audiovisual electrodes where nouns versus adjectives was the only significant predictor in the GLM analysis, 6 electrodes (46%, 4 in the left-LOF, and 2 in left superior temporal gyrus) robustly distinguished nouns and adjectives matched for their frequency of occurrence, like the example electrode in **Figures 2** and **S5** whereas the other electrodes maintained their statistically significant selectivity in most but not all conditions (note that the statistical power is lower in the matched analyses due to subsampling).

We considered the possibility that frequency of occurrence and word surprisal may also contribute to neural responses. There was a significant negative correlation between frequency of occurrence and word surprisal (calculated using GPT-2, **Methods**) for both word1 (r=-0.62, p<0.001, two-tailed t test) and word2 (r=-0.39, p<0.001, two-tailed t test). Thus, we made separate GLMs for each of these predictors given that GLMs are less effective with correlated variables. **Figure S5e,f** shows GLMs for the example electrode in **Figure 2a-d**, separately for frequency of occurrence (**Figure S5e**) and GPT-2 surprisal (**Figure S5f**). Part of speech was the only GLM predictor that significantly explained the neural responses of this electrode for both variables. A majority of electrodes from **Figure 2l** had part-of-speech as the only significant contributor for GLMs with frequency of occurrence (n=8 with POS as the only significant predictor; n=5 electrodes ceased to have any significant predictor, instead of an alternative predictor), and for word surprisal (n=11). **Figure S5g** shows the distribution of different variables that co-explained neural variability along with the parts of speech for the GLM with surprisal. **Figure S5h** shows their brain locations.

There were two subcategories of nouns and two subcategories of adjectives within our stimulus set: animals/food, and concrete/abstract (**Table S3**). We asked whether the electrodes that showed differential responses generalized across different word subcategories. The example electrode in **Figure 2a-i** did not show differences between the two noun or adjective subcategories for either auditory stimuli (**Figure S7a, b, f, g**), visual stimuli (**Figure S7c, d, h, i**), word 1 (**Figure S7a, c, f, h**), or word 2 (**Figure S7b, d, g, i**). Of the 13 audiovisual electrodes where noun versus adjective was the only significant predictor in the GLM analysis, 8 electrodes (62%) showed generalization across different noun or adjective subcategories. The remaining 5 electrodes (38%) showed a statistically significant difference between the two noun subcategories or between the two adjective subcategories (Table S5; note that the statistical power is also lower in this analysis, which includes approximately half the number of trials). **Figure S7j-r** shows one of the exceptions—an electrode in the left LOF that showed a significant response only for food nouns. This selectivity was particularly pronounced for visual stimuli (**Figure S7l, m, q, r**), but was also apparent for auditory stimuli (**Figure S7j, k, o, p**), and was evident for both word1 and word2. In sum, differences in selective responses to nouns versus adjectives were particularly prominent and clustered in the left lateral orbitofrontal cortex, persisted across different word lengths, whether the word was used in a grammatically correct phrase or not, after equalizing word occurrence frequency, and generalized across different noun or adjective subcategories.

### Neural signals enhanced for nouns versus adjectives were anatomically segregated

Of those electrodes uniquely selective for part of speech, 79% showed responses that were significantly stronger for nouns compared to adjectives (β_NvsA_ > 0) as illustrated by the example in **Figure 2a-i**. The remaining 21% showed responses that were stronger for adjectives compared to nouns (β_NvsA_ < 0) as illustrated by the example in **Figure S6j-s** (**Table S5)**. There were also marked differences in the dynamics underlying the responses to nouns and adjectives for auditory stimuli, but not for visual stimuli. For auditory stimuli, the difference in the onset time between nouns and adjectives was larger for noun-preferring electrodes (550 ± 107 ms) than for adjective-preferring electrodes (312 ± 94 ms, ranksum test, p<0.05). For visual stimuli, the difference in the onset time between nouns and adjectives was not different between noun-preferring electrodes (425 ± 107 ms) and adjective-preferring electrodes (437 ± 134 ms, ranksum test, p>0.05). There was a significant correlation between auditory and visual difference onset times for noun-preferring electrodes (Pearson R = 0.80, p<0.01, n=10) but not for adjective-preferring electrodes (Pearson R = −0.70, p>0.05, n=3).

When we displayed the electrode locations on the brain, we observed an anatomical separation between these two groups of responses (**Figure 2m,n**, x-axis: lateral to medial, y-axis: anterior to posterior, z-axis: ventral to dorsal). We compared noun- versus adjective- preferring electrodes along 3 axes of Montreal Neurological Institute 305 Coordinates (MNI305, units abbreviated as m.u.)^44^. Along the medial to lateral axis (x-axis in **Figure 2m,n**, zero being more medial), noun-preferring electrodes had a mean of 25.3±6.2 m.u. and adjective-preferring electrodes had a mean of 47.3±7.7 m.u. (p<0.01, ranksum test). Along the ventral-dorsal axis (z-axis in **Figure 2m**), noun electrodes had a mean of −12.17±5.3 m.u. and adjective electrodes had a mean of −3.7±1.7 m.u. (p<0.05, ranksum test). Along the posterior-anterior axis (y-axis in **Figure 2n**), noun electrodes had a mean of 21.4±18.9 m.u. and adjective electrodes had a mean of − 2.7±25.8 m.u. (p<0.05, ranksum test). **Table S6** summarizes the locations of noun- vs adjective-preferring electrodes across brain regions. A permutation test for these electrodes showed that electrodes in the LOF tended to show stronger responses to nouns (∼90% β_NvsA_ > 0, p<10^-4^, permutation test, n=10^6^ iterations, **Methods**).

### A population of electrodes in the lateral orbitofrontal cortex can distinguish nouns from adjectives in individual trials and generalizes across word positions and modalities

The raster plots in **Figure 2e-h** and **Figures S4-S7** qualitatively show that the responses are strong enough to be discerned in individual trials. To assess whether information about part of speech was available in individual trials, we used a machine learning pseudopopulation approach by combining electrodes within anatomically defined brain regions in the Desikan-Killiany Atlas^45^. We binned the responses in 100 ms time bins and used the top-N principal components that explained more than 70% of the variance in the training data for all the electrodes. We trained an SVM classifier with a linear kernel to distinguish between nouns and adjectives and tested the classifier on held-out data (**Methods**). In Figure 3, the same word could randomly be part of the training and test set while still using independent train and test data across trials or word positions. **Figure 3** shows decoding accuracy for the left (**Figure 3a,d,g**) and the right (**Figure 3b,e,h**) LOF as a function of time from word onset. When trained using data from both word1 and word2 with combined auditory and visual features, there was a statistically significant decoding performance starting approximately at 300 ms after word onset and reaching a peak of 63.6 ± 1.1% at ∼500 ms after word onset in the left LOF (**Figure 3a**). Statistical significance was assessed by comparing with a control where noun and adjective labels were randomly shuffled (**Methods**). Even though there were almost twice as many electrodes in the right LOF compared to the left LOF (**Figure 1b-g, Table S2**), decoding performance was higher for the left LOF compared to the right LOF (compare **Figure 3a** versus **Figure 3b**). The differences between the left and right LOF persisted after randomly subsampling to equalize the number of electrodes across hemispheres for all regions (**Figure 4a,b**).

**Fig. 3.**
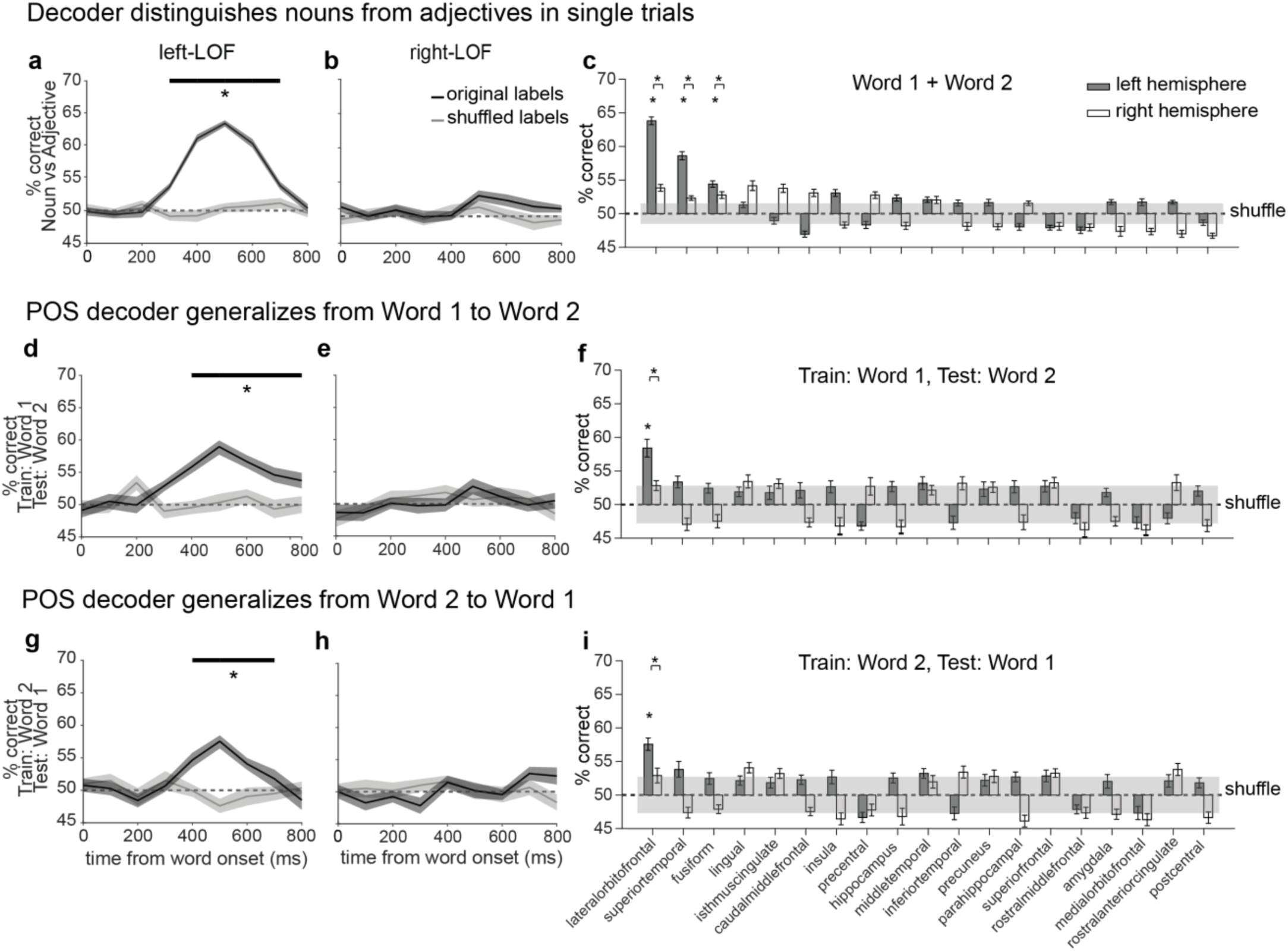
Neural signals distinguishing nouns and adjectives in single trials generalize across Word1, and Word2. **a, b, d, e, g, h.** Average cross-validated performance of a support vector machine classifier (SVM, 80% training/20% test) decoding nouns versus adjectives for all electrodes in the left lateral orbitofrontal cortex (LOF) **(a, d, g)** or the right LOF **(b, e, h)**. The dotted horizontal black line shows the chance level. Shaded areas denote s.e.m. Solid horizontal black bar shows time points where performance significantly differed from chance (100 random shuffles, ranksum test, p<0.01). The inputs to the SVM included the top-N principal components of the electrode response that explained >70% variance for the training data at each time bin (**Methods**). **a, b**: Features from auditory and visual responses were combined and used for training and testing on a dataset of both Word1 and Word2 trials. **d,e**: Generalization across word order was evaluated on a dataset where Word1 trials were used for training and word2 trials were used for testing. **g, h**: Training on Word2 and testing on Word1. Black: original labels; Gray: shuffled labels. (see **Figure 4** for decoding performance when the number of electrodes was the same across all regions and both hemispheres) **c, f, i.** Summary of average of max-decoding performance for distinguishing nouns versus adjectives in each hemisphere (dark: left; white: right) for different brain regions. Bottom asterisks denote regions with significant decoding performance with respect to chance and performance from the real and null distribution do not overlap within 3 standard deviations of each other (p<0.01, ranksum test, corrected for multiple comparisons, **Methods**). Shaded box: maximum of the mean ± SD. for the null distribution across all regions. Top asterisks with a U-bracket denote significant differences between decoding accuracy of the left versus the right hemisphere (p<0.01, ranksum test, corrected for multiple comparisons). Regions are sorted in descending order of performance in panel **c**. **c**: Classifiers were trained and tested with features from both Word1 and Word2 trials. **f**: Classifiers were trained on Word1 trials and tested on Word2 trials. **i**: Classifiers were trained on Word2 trials and tested on Word1 trials. (see **Figure 4** for controls on word1-only, word2-only, audio-only, visual-only, audio-to-vision and vision-to-audio performance.)

**Fig. 4.**
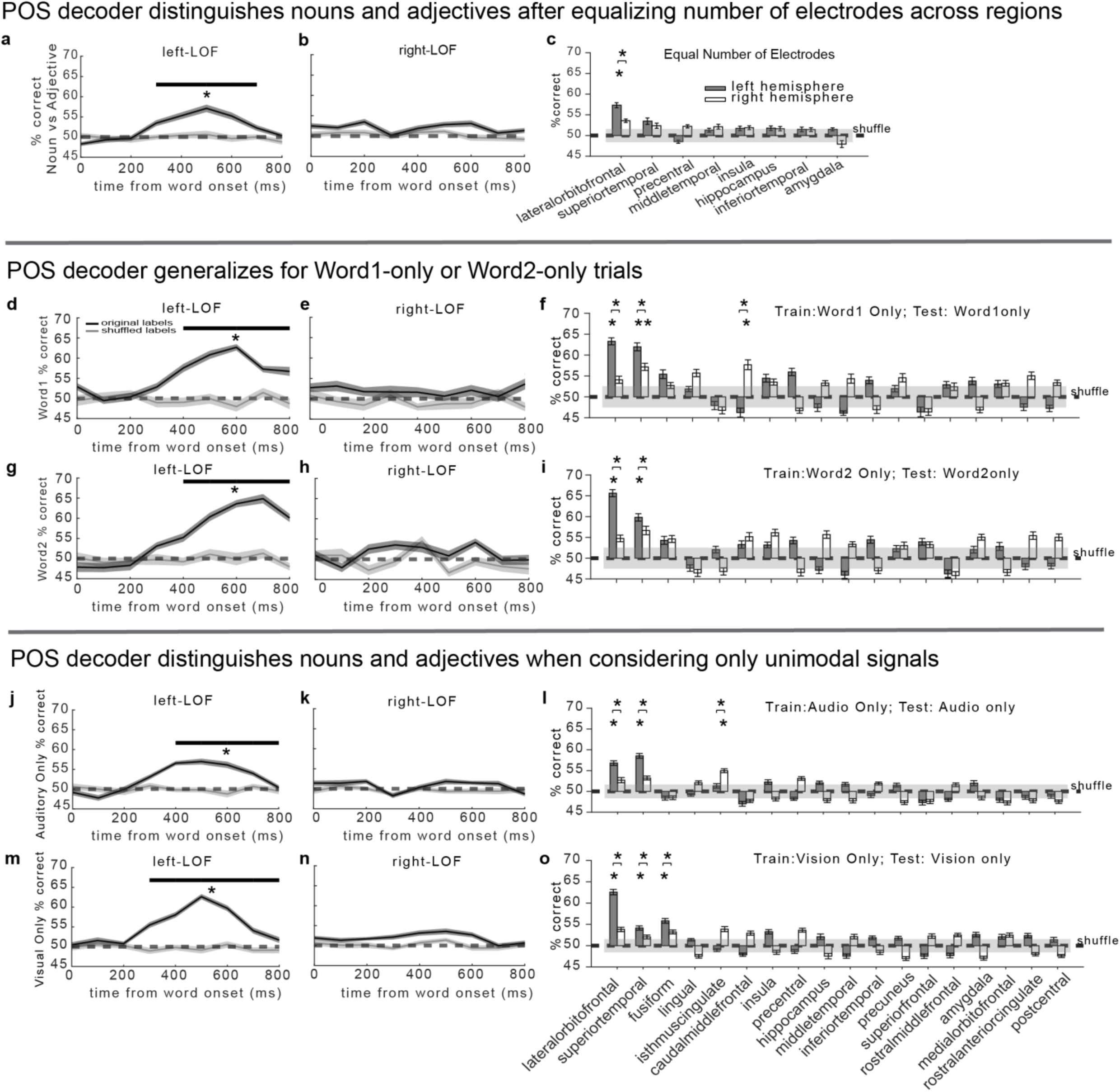
Neural signals from left-LOF distinguish nouns and adjectives for i) regions normalized for electrode numbers, ii) given word position in phrase and iii) given modality. **a-b.** Average cross-validated performance of a support vector machine classifier (SVM, 80% training/20% test) decoding nouns versus adjectives for 8 randomly subsampled electrodes in the left lateral orbitofrontal cortex (LOF) (**a**), and in the right LOF (**b**). Black: original labels; Gray: shuffled labels. The dotted horizontal black line shows the chance level. Solid horizontal gray bar shows time points where decoding from correct labels significantly differed from that of shuffled labels (100 random shuffles of the data, ranksum test, p<0.01). The inputs to the SVM were 100 ms time bins from word onset containing the top-N principal components of the electrode response at each bin that explained >70% variance for the training data (**Methods**). **c.** Summary of average of max-decoding performance for distinguishing nouns versus adjectives across both hemispheres (left hemisphere: dark gray bars; right hemisphere: white bars) for different brain regions when a total of 8 electrodes was taken from each hemisphere in each region for the decoding. Regions with less than 8 electrodes in either hemisphere were omitted. Asterisk: significant hemisphere within a Desikan-Killiany defined brain region (p<0.01, ranksum test, corrected for multiple comparisons, and performance from the real and null distribution do not overlap within 3 standard deviations of each other) (**Methods**). Gray box: maximum mean ± s.t.d. for the null distribution across all regions. Asterisk with a U-bracket: significant difference between decoding accuracy of the left versus the right hemisphere (p<0.01, ranksum test, corrected for multiple comparisons). Regions are sorted in descending order of performance in panel **c**. **d, e, g, h, j, k, m, n.** Average cross-validated performance of a support vector machine classifier (SVM, 80% training/20% test) decoding nouns versus adjectives for all electrodes in the left lateral orbitofrontal cortex (LOF) (**d,g,j,m**), and in the right LOF (**e,h,k,n**). The dotted horizontal black line shows the chance level. Shaded areas denote s.e.m. Solid horizontal black bar shows time points where performance significantly differed from chance (100 random shuffles, ranksum test, p<0.01). The inputs to the SVM included the top-N principal components of the electrode response that explained >70% variance for the training data at each time bin (**Methods**). Features from auditory and visual responses were combined and used for training and testing on datasets of word1 (**d,e**) and word2 trials (**g,h**). Using a combined dataset of word1 and word2 trials, the decoding performance was evaluated for audio-only (**j,k**) and vision-only features (**m,n**). **f, i, l, o.** Max-decoding accuracy across regions (left: dark; right: white) for the same conditions. Bottom asterisks: significant above chance; top U-bracket asterisks: left–right differences (p<0.01, corrected).

In **Figure 3a-c**, word 1 and word 2 were combined. Decoding performance in the left LOF was also significant when separately considering word 1 (**Figure 4d-f**) or word 2 (**Figure 4g-i**). Furthermore, the machine learning classifier was able to generalize across word positions, as evidenced by the decoding performance when training on the responses to word 1 and testing on the responses to word 2 (**Figure 3d-f**), and vice versa (**Figure 3g-i**). Similarly, auditory and visual trials were combined in **Figure 3a-c**. Decoding performance in the left LOF was also significant when separately considering auditory trials (**Figure 4j-l**) and visual trials (**Figure 4m-o**). The machine learning classifier was also able to generalize across modalities, as evidenced by the decoding performance when training on auditory trials and testing on vision trials (**Figure 5a-c**) and vice versa (**Figure 5d-f**). Furthermore, the classifier also generalized when we trained on given subclasses of nouns and adjectives (e.g., food nouns versus abstract adjectives), and tested with different subclasses (e.g., animal nouns versus concrete adjectives) (**Figure 5g-i**). To investigate the contribution of multimodal sites in decoding, we removed the multimodal responsive electrodes (**Table S2**) from the left LOF before performing the decoding analyses. After the removal of the multimodal responsive electrodes, the performance of the decoder using left LOF was not different from chance (51.9±3.4%, ranksum test, p>0.05). However, when we exclusively used multimodal electrodes, the performance was significantly greater than chance (63.8±2.6%, ranksum test, p<0.01). In sum, the encoding of part-of-speech information generalized across **word position** and **sensory modality**.

**Fig. 5.**
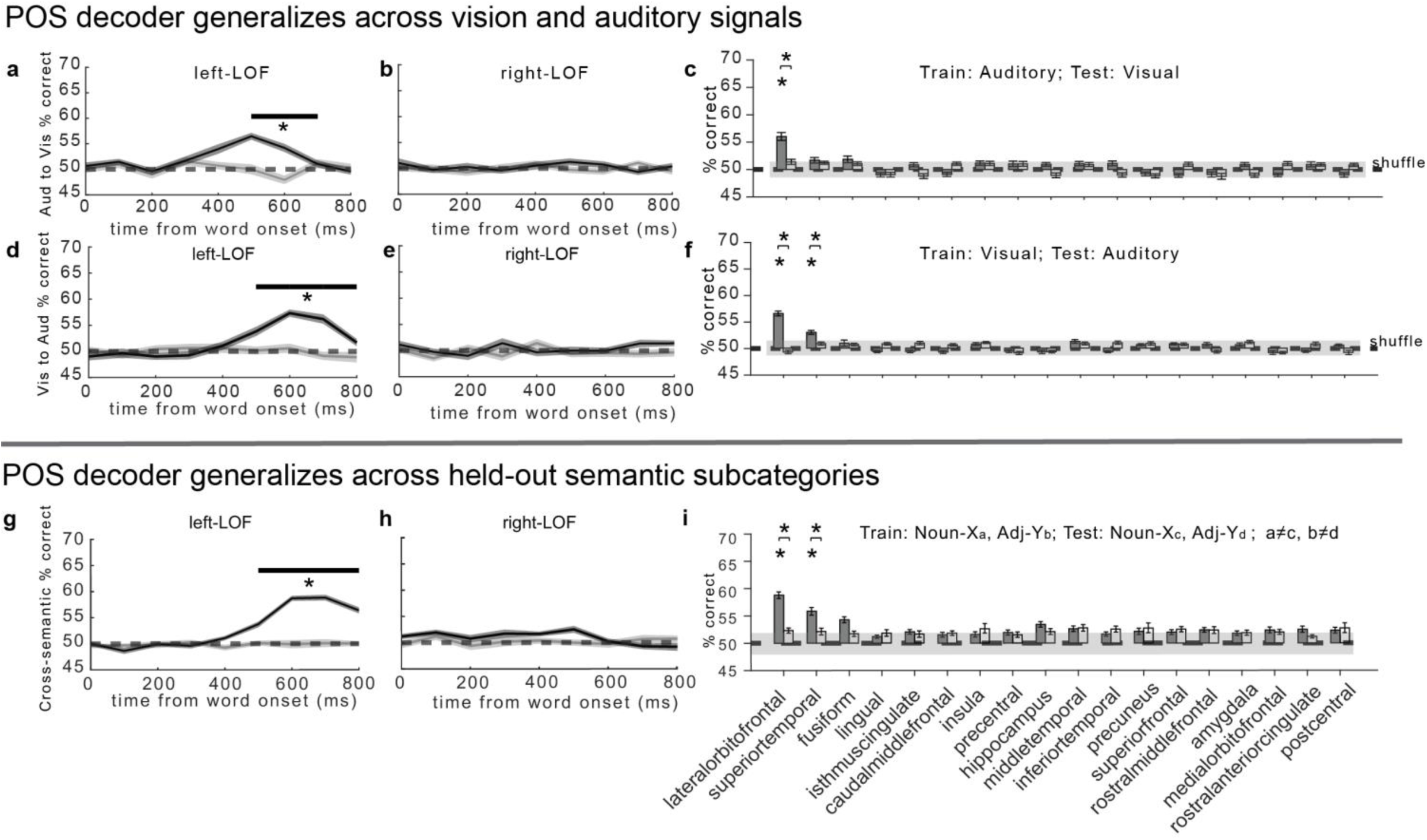
Neural signals from left-LOF distinguish nouns and adjectives i) across modalities, and ii) word subclasses. **a, b, d, e.** Average cross-validated performance of a support vector machine classifier (SVM, 80% training/20% test) decoding nouns versus adjectives for all electrodes in the left LOF (**a, d**) and the right LOF (**b, e**). Using a combined dataset of word1 and word2 trials, the classifier was trained on auditory features and tested on visual features (**a, b**), and trained on visual features and tested on auditory features (**d, e**). SVM inputs were the top-N principal components of the electrode response that explained >70% variance for the training data at each time bin (Methods). The dotted horizontal black line shows the chance level. Shaded areas denote s.e.m. Solid horizontal black bar shows time points where performance significantly differed from chance (100 random shuffles, ranksum test, p<0.01). **c, f.** Max-decoding accuracy across regions (left: dark; right: white) for the audio (train)→vision (test) (**c**) and vision→audio (**f**) generalization conditions. Bottom asterisks: significant above chance; top U-bracket asterisks: left–right differences (p<0.01, corrected). **g, h.** Average cross-validated performance of an SVM in the left (**g**) and right LOF (**h**), trained on Noun-Xa versus Adjective-Yb and tested on Noun-Xc versus Adjective-Yd, where {Xa, Xc} and {Yb, Yd} are different subclasses of nouns and adjectives, respectively (e.g., Xa = animal nouns, Xc = food nouns). This yields 4 combinations for testing noun-versus-adjective decoding across semantic subcategories; the average across these 4 tests for 100 random shuffles is plotted. Solid horizontal black bar shows time points where performance significantly differed from chance (100 random shuffles, ranksum test, p<0.01). **i.** Summary of average max-decoding performance for distinguishing nouns versus adjectives across subclasses in each hemisphere (dark: left; white: right) for different brain regions. Bottom asterisks: significant above chance; top U-bracket asterisks: left–right differences (p<0.01, corrected).

We extended the analyses in **Figure 3a,b,d,e,g,h** to all other regions in the Desikan-Killiany atlas. In addition to the left LOF, the left superior temporal cortex and the left fusiform cortex also showed statistically significant decoding performance (**Figure 3c**). However, in contrast to the results for the left LOF, the decoding results for other regions were less robust (**Figure 4c**) and did not consistently generalize across word position (**Figure 3f**,**i**) or across modalities (**Figure 5 c,f**).

### Multimodal neural signals distinguishing different parts of speech are conserved across languages

One of the participants was fluent in two languages, English and Spanish. Therefore, this patient provided an opportunity to ask whether the neural signals discriminating between different parts of speech were language-specific or showed invariance across languages. All the words were translated into Spanish by a native Spanish speaker, and the task was repeated in both languages. **Figure 6a-h** shows the responses of an example electrode located in the left LOF (**Figure 6k**). This electrode showed a stronger response to nouns compared to adjectives for auditory stimuli (**Figure 6a**, **b**, **e**, **f**), for visual stimuli (**Figure 6c**, **d**, **g**, **h**), for Word 1 (**Figure 6a**, **c**, **e**, **g**), and for Word 2 (**Figure 6b**, **d**, **f**, **h**). Interestingly, the separation between nouns and adjectives was evident both when the words were presented in English (**Figure 6a-d**) and when the words were presented in Spanish (**Figure 6e-h**). The GLM analysis showed that nouns versus adjectives was the only significant predictor in English trials (**Figure 6i**), and Spanish trials (**Figure 6j**). All in all, there were three electrodes in this participant that showed a multimodal response selective for part of speech. We illustrate the effect with one example electrode; the other two LOF electrodes showed qualitatively similar POS selectivity. All three of these electrodes were in the left orbital H-shaped sulcus within the LOF (**Figure 6k**, green).

**Fig. 6.**
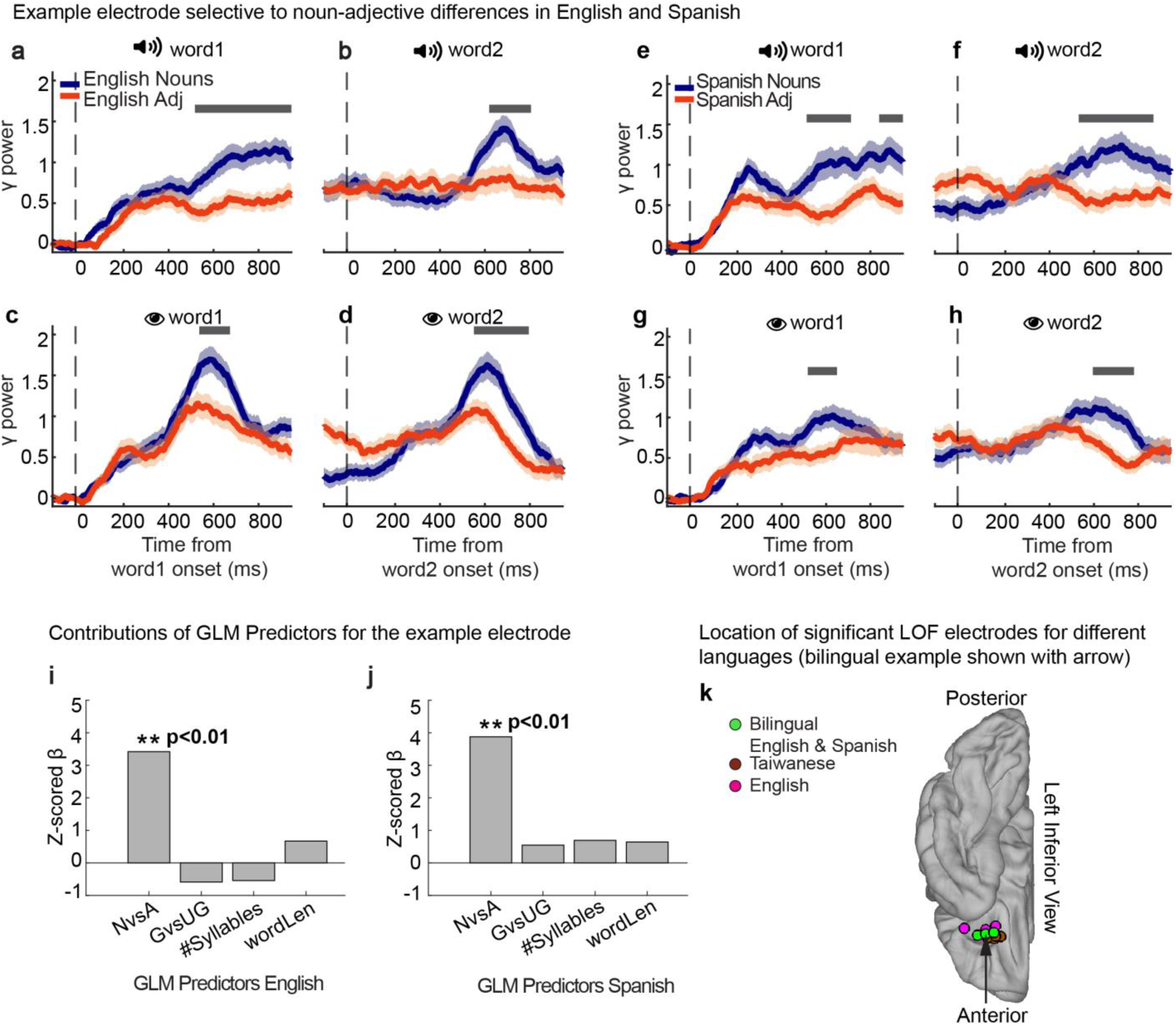
Neural signals in left LOF generalize across languages in a bilingual subject and in monolingual subjects. **a-h.** Trial averaged responses of an electrode in the left lateral orbitofrontal cortex from a bilingual patient. The format follows **Figure 2a-d**. (**a-d**) English words (audio: n=190 grammatical and 185 ungrammatical trials; vision: n=189 grammatical and 191 ungrammatical trials). (**e-h**) Spanish words (audio: n=184 grammatical and 184 ungrammatical trials; vision: 184 grammatical and 186 ungrammatical trials). Auditory responses (**a, b, e, f**). Visual responses (**c, d, g, h**). Word 1 (**a, c, e, g**) and Word 2 (**b, d, f, h**). **i,j.** Z-scored β coefficients for Generalized Linear Model to predict area under the curve (AUC) for the English experiment (**i**) and for the Spanish experiment (**j**). The AUC computed between 200 ms and 800 ms post word onset using four task predictors: Noun versus Adjectives, Grammatical versus Ungrammatical, number of syllables (auditory presentation) and word length (visual presentation). Asterisks denote statistically significant coefficients corrected for multiple comparisons **(Methods)**. The word order for grammatically correct trials in English is an adjective followed by a noun, such as “green apple”. This word order gets flipped in grammatically correct Spanish trials. **k.** Inferior view of all the 9 out of 38 electrodes (8 audiovisual: **Figure 2l,** 1 visual-only: **Figure S4u,v,** see **Table S4** and **S7**) in the left lateral orbitofrontal cortex that showed noun versus adjective differences across different languages in which the experiment was conducted (significant Nouns versus Adjectives β, p<0.01 corrected for multiple comparisons). These electrodes come from 4 different subjects. Electrodes from the bilingual patient are in green with a black arrow indicating the example electrode. Electrodes from one monolingual English patient are in pink and those from 2 monolingual Taiwanese patients are in brown.

In addition to this bilingual participant, the task was run in monolingual participants who spoke English (n=16 participants) and monolingual participants who spoke the Taiwanese dialect of Chinese (hereafter referred to as Taiwanese, unless otherwise specified, to avoid confusion regarding pronunciation and writing differences, n=3 participants, **Table S1**). In **Figure 6k**, we show all electrodes from the left LOF that showed part-of-speech encoding from different participants (**Table S7**). LOF electrodes in 4 out of 8 participants (50%) with coverage in the region exhibited differences between nouns and adjectives. We also indicate the language in which this difference was observed whether it be English (pink), Taiwanese (brown) or bilingual English/Spanish (green). Electrodes separating parts of speech from monolingual participants were also clustered in the same region.

In sum, electrodes in the left LOF robustly distinguished between parts of speech for both auditory and visual presentations of stimuli across participants speaking English, Taiwanese or Spanish. Additionally, in one bilingual subject, the same electrodes showed robust part-of-speech selectivity that generalized across English and Spanish.

## Discussion

We described neurophysiological signals that selectively discriminate between two parts of speech, nouns and adjectives (**Figure 2**). This selectivity was robust to orthographic variables such as word length, phonetic features such as number of syllables, and word occurrence statistics (**Figure 2**). This selectivity for part of speech generalized across sensory modalities (**Figures 2**, **4, 5**), word positions, grammatical correctness and motor outputs (**Figures 2**, **3**, **4, 5, 6**), and semantic groups of nouns and adjectives (**Figure S7**). These neurophysiological signals enabled discrimination between parts of speech even in single trials (**Figures 2, 3, 4, 5**). Electrodes that uniquely distinguished nouns from adjectives were clustered within a small, circumscribed region of the lateral orbitofrontal cortex, lateralized to the left hemisphere (**Figure 2**, **6**). Neural discrimination of nouns from adjectives was apparent in the LOF in English-speaking and Taiwanese-speaking participants (**Figure 6**). Interestingly, in a bilingual participant, the same electrodes within the left LOF distinguished nouns and adjectives in both English and in Spanish (**Figure 6**).

We identify a highly circumscribed region in the posterior left lateral orbitofrontal cortex (posterior H-shaped sulcus) whose activity reliably distinguishes nouns from adjectives in a manner that generalizes across sensory modality, word position, semantic subclass, and languages. This localization is not predicted by classic perisylvian accounts that emphasize inferior frontal and temporal cortices for grammatical and lexical operations. These results provide an anatomically and temporally resolved neural target for testing how grammatical category signals interact with canonical language regions during phrase processing and comprehension.

Clinically, a focal LOF locus carrying invariant part-of-speech signals may inform intracranial language mapping and interpretation of language risk when resections approach posterior LOF. A critical goal of cortical resections in epilepsy patients is to cure seizures without interfering with cognitive function. As such, given the strong lateralization and ubiquitous role for language operations in cognition, it is extremely important to precisely understand the neural structures that support language in these patients; the current results could help guide surgical approaches for epilepsy.

In English and other languages, some words can be used both as a noun and as an adjective (e.g., a *light* color versus turn on the *light*). In most instances, one usage is more frequent than the other. For this study, we selected nouns and adjectives that are highly overrepresented in their labeled part of speech. The nouns and adjectives in this study were used in their labeled part-of-speech category at far higher frequency than in any alternative part-of-speech category, as quantified by the occurrence ratios in **Table S8**. Similarly, some words can be used both as a noun and as a verb (e.g., “long *race”* versus “*race* you to the top”); all the nouns in this study are highly overrepresented in their usage as nouns versus verbs (**Table S8**). Thus, the words used in this study had a prototypical interpretation as either, noun or adjective. The distinction between POS includes their grammatical roles but also their associated semantic connotations (e.g., nouns typically refer to things and adjectives to the attributes of those things).

In languages like English, nouns and adjectives follow a specific grammatical order (i.e., adjectives precede nouns). Other languages reverse this order. In Spanish, adjectives typically follow nouns, though the English order can also be used. It is thus interesting to observe that many electrodes demonstrated strong selectivity for nouns versus adjectives, irrespective of their position within the two-word phrases. Furthermore, in one bilingual participant, the neural responses separated nouns and adjectives in both languages despite the fact that the grammatical order is typically reversed between English and Spanish. This bilingual experiment emphasizes the role of part-of-speech: since the grammatical structure reverses between English and Spanish while noun-preferring responses remain robust, the effects cannot be explained solely by word order, first-word surprisal, or word frequency. However, it is conceivable that the strong part-of-speech selectivity independent of grammar shown here could be linked to the two-word phrase structures. The underlying reason for the dissociation between nouns and adjectives could also be either semantic (adjectives refer to properties of things, nouns to individual things), or syntactic (e.g., head vs adjuncts). The adjective-noun two-word phrases cannot dissociate part-of-speech from head-vs-adjunct status because the noun is always the phrasal head and the adjective always the modifier. Another possibility is that the representation of nouns versus adjectives is invariant to grammatical usage rules. Non-invasive scalp electroencephalography and magnetoencephalography signals have revealed correlates of language processing with a wide range of onset times from approximately 100 ms all the way to well over 600 ms (for a review, see^46^). The earliest onset signals commencing between 100 and 300 ms after stimulus onset, sometimes referred to as early left anterior negativity, have been associated with grammatical violations, but previous studies have not documented any invariance in the representation of parts of speech and there is disagreement about whether these early signals are even associated with language^46^. Our work reports an invariant distinction between nouns and adjectives in the LOF commencing at approximately 400 ms after stimulus onset, which is consistent with part-of-speech being represented well after the onset of modality-specific purely visual and auditory signals.

A remarkable hallmark of language is its universality. We can interpret the word *cat* when uttering the word, writing it, listening to it, reading it, and even when examining a photograph of a cat. It is therefore tempting to speculate that there may be an invariant representation of language concepts in the brain. Several studies have examined putative correlates of language processing using only unimodal signals (e.g.^11–15,18,24,29,34,35^). While we observed electrodes that distinguished between parts of speech only in the auditory stimuli or only in the visual stimuli, the responses of those electrodes could be partly explained by other variables including number of syllables, word frequency, or grammar. Using strict criteria and after controlling for confounding variables, most electrodes that distinguished nouns from adjectives showed selectivity during both auditory and visual presentation. Future work should evaluate whether the same electrodes also distinguish parts of speech when participants utter words, write them, or when examining images. An intriguing study described neurons in the human medial temporal lobe that respond selectively to images and their corresponding text and sound descriptions^47–48^. However, these medial temporal lobe neurons do not seem to distinguish between different parts of speech and their responses seem to be connected with the formation of memories rather than the internal representation of language^49^. Indeed, there exist strong anatomical and functional connections between the medial temporal lobe and frontal regions that could link language and memory formation^50^.

The lateral orbitofrontal (LOF) cortex constitutes a large expanse of neocortex within the frontal lobe, spanning Brodmann areas (BA) 10, 11, 12 (called BA47 in humans due to cytoarchitectural differences from monkeys) and 13^51–53^. Neuroanatomical tracings from rats, mice, and macaques have identified LOF as a nexus of many inputs^53^ conveying olfactory, gustatory, visual, auditory, somatosensory, and visceral-sensory information. The LOF has been associated with many cognitive functions, including multisensory integration, working memory, long-term memory consolidation, reward processing, social interactions, decision making, and emotion processing^50,52,54–58^. This heterogeneity might be partly ascribed to investigations probing different cognitive tasks, as in the case of the proverbial blind men sampling different parts of an elephant. Given the prominent role of language in cognition, it is conceivable that previous studies that describe other roles of the LOF did not probe its possible associations with language. However, it is even more likely that descriptors like LOF that refer to such large brain areas would inevitably fail to uncover specific functionality. The current results point to a rather well circumscribed location within LOF, the posterior part of the H-shaped sulcus in the left hemisphere. In humans, this location overlaps with BA13-lateral and BA47-medial and has been shown to have a strong convergence of auditory and visual inputs^52,53,59^. Interestingly, work on Primary Progressive Aphasia, and frontotemporal lesions implicate the orbitofrontal cortex in word and sentence comprehension deficits^7, 59–61^ (see also^28^). In these studies, the orbitofrontal cortex, dorsal premotor cortex, temporoparietal junction (canonical Wernicke’s area), and pars opercularis were associated with sentence comprehension and grammatical production aphasias (evaluated with complex grammatical output requiring planning and motor production). Word comprehension and naming deficits were assessed using binary perceptual choice tasks, implicating the orbitofrontal cortex and the anterior temporal lobe (ATL). Consistent with extensive work documenting the lateralization of language functions, the results presented here also show a strong predominance of the left hemisphere in the representation of part of speech, despite the fact that there were more electrodes sampling signals from the right hemisphere.

Several limitations in the current work are worth noting. First, all the results reported here are derived from patients with epilepsy. The invasive study of epilepsy patients constitutes the predominant way to access neurophysiological signals from the human brain^62,63^. Neurophysiological studies in other patient populations (e.g., paraplegic patients, Parkinson’s patients, brain tumor patients) typically target specific regions that are not known to be associated with language processing. Caution should be exercised in the interpretation of results from patient populations. To the best of our knowledge, all patients used language fluently and had no language impediments, but one should be aware of the possibility that epilepsy could potentially impact the representation of language. Second, the electrode locations are strictly dictated by clinical criteria. Our sampling of brain activity is extensive but not exhaustive (**Figure 1**, **Tables S1-S2**). It is quite possible that other areas not examined here may also reveal neural correlates of parts of speech and that the regions we found interact with other relevant brain areas. As shown in **Table S7**, the POS selective electrodes were present in approximately 50% of participants with LOF coverage. While the participants showed a wide age range, individual differences in electrode placement, rather than age, likely account for variability in observed effects. Additionally, the limitations in the number of participants and number of electrodes, especially a single bilingual participant, should be considered when interpreting the results. Future studies with more electrode coverage and more participants would be welcome. Third, the current work focuses on two parts of speech. Nouns and adjectives do *not* constitute an exhaustive list of parts of speech and future work should examine the representation of pronouns^64^, verbs, adverbs, prepositions, and conjunctions. Finally, our work provides a *correlate* of the representation of POS, but future work should evaluate whether any such signals are *causally* required for online language interpretation. Establishing a causal role will require perturbation approaches (e.g., electrical stimulation of LOF during tasks that manipulate grammatical category and composition or lesion–symptom analyses that test whether disrupting LOF selectively impairs POS-based judgments).

These results provide initial glimpses into highly localized structures that represent a fundamental component of language that has been extensively studied by linguists for decades, the functional role of different words within a sentence. The representation of nouns versus adjectives in the human brain is invariant to the presentation modality, word properties, grammar, and the noun and adjective subcategories tested here. Furthermore, the representation even generalizes across different languages. These findings open interesting questions about how next-token prediction in language models relates to the brain’s implicit biases that give rise to stable, invariant representations of parts of speech. These observations open the doors to begin to elucidate the neural representation of more complex language concepts and to bridge the extensive work in language and linguistics to their underlying neural representations.

## RESOURCE AVAILABILITY

### Lead Contact

Further information and requests for resources should be directed to and will be fulfilled by the Lead Contact, **Gabriel Kreiman**.

### Materials Availability

All the stimuli are publicly available at the repository listed below.

### Data and Code Availability

All data and code are publicly available at the following site: https://kreimanlab.com/code/neural-coding-for-language-representations-in-the-human-brain/ The pseudocode is included in the accompanying *Readme.docx* file in the same repository.

## ACKNOWLEDGMENTS

We thank Marcelo Armendariz and Yen-Ling Kuo for helping translate the task to Spanish and Taiwanese, respectively. We thank Professors Alfonso Caramazza and Boris Katz for suggestions and insightful comments throughout this work.

## AUTHOR CONTRIBUTIONS

The experiment was designed by PM and GK. All the surgeries were performed by YS, HY, JRM, and SS. DW helped coordinate all the experiments. All the data were collected and analyzed by PM. The manuscript was written by PM and GK with feedback from all the authors.

## SUPPLEMENTAL INFORMATION

Supplement PDF containing **Tables S1–S8** and **Figures S1–S7**

## STAR★Methods

### Key Resource Table

| REAGENT or RESOURCE | SOURCE | IDENTIFIER |
| --- | --- | --- |
| **Deposited data** |  |  |
| Neurophysiological recordings, stimuli, and analysis code | This paper | https://kreimanlab.com/code/neural-coding-for-language-representations-in-the-human-brain/ |
| **Experimental models: Human participants** |  |  |
| 20 neurosurgical patients undergoing sEEG monitoring | Collaborating Hospitals | IRB approved; see Table S1 |
| **Software and algorithms** |  |  |
| MATLAB | MathWorks, Natick, MA | https://www.mathworks.com/; RRID:SCR_001622 |
| Psychtoolbox-3 | Brainard, 1997; Pelli, 1997 | http://psychtoolbox.org/; RRID:SCR_002881 |
| FreeSurfer | Fischl, 2012 | https://surfer.nmr.mgh.harvard.edu/; RRID:SCR_001847 |
| iELVis | Groppe et al., 2017 | https://github.com/iELVis/iELVis |
| GPT-2 | Radford et al., 2019 | https://github.com/openai/gpt-2 |
| **Other** |  |  |
| XLTEK neurophysiological recording system | Natus Medical, Oakville, ON, Canada | N/A |
| Ad-Tech sEEG depth electrodes | Ad-Tech, Racine, WI, USA | N/A |

### Experimental Model and Subject Details

#### Preregistration

This study was preregistered on the Open Science Framework (OSF) website. The preregistration DOI is: https://doi.org/10.17605/OSF.IO/8TU2G.

#### Participants

We recorded data from 20 participants (9 male, 9-53 years old, 2 left-handed, 2 ambidextrous, **Table S1**) with pharmacologically resistant epilepsy (**Figure 1a**). All experiments were conducted while participants stayed at Children’s Hospital Boston (CHB), Brigham and Women’s Hospital (BWH), or Taipei Veterans General Hospital (TVGH). All studies were approved by each hospital’s institutional review boards and were carried out with the participants’ informed consent.

#### Recordings and Electrode Locations

Participants were implanted with intracranial electrodes via stereo electroencephalography (sEEG) (Ad-Tech, Racine, WI, USA). Neurophysiological data were recorded using XLTEK (Oakville, ON, Canada), Bio-Logic (Knoxville, TN, USA), Nihon Kohden (Tokyo, Japan), and Natus (Pleasanton, CA). The sampling rate was 2048 Hz at BCH and TVGH, and 1024 Hz or 512 Hz at BWH. All data were referenced in a bipolar montage. There were no seizure events in any of the sessions. Electrode locations were decided based on clinical criteria for each participant. Electrodes in the epileptogenic foci, as well as pathological areas, were removed from analyses. The total number of electrodes after bipolar referencing and removing electrodes with no signal, line noise or recording artifacts was 1,801^65^.

Following implantation, electrodes were localized by co-registration of pre-operative T1 MRI and post-operative CT scans using the iELVis software^44^. We used FreeSurfer to segment MRI images, upon which post implant CT was rigidly registered^66^. Electrodes were marked in the CT aligned to pre-operative MRI using the Bioimage Suite^67^. The Desikan-Killiany (DK) atlas was used to assign the electrodes locations. **Figure 1b-g** and **Table S2** show the locations of all the electrodes.

#### Experiment Design

All visual stimuli were displayed on a 15.4 inch 2,880 × 1,800 pixel LCD screen using the Psychtoolbox in MATLAB (Natick, MA) and a MacBook Pro laptop (Cupertino, CA). The stimuli were positioned at eye level at about 80 cm from the participant and each word subtended approximately 3 degrees of visual angle. Sounds were played from the speakers of a MacBook Pro 15.4 at 80% loudness using the Psychtoolbox in MATLAB^68^. We used the USB-1208FS-Plus device from Measurement Computing Corporation (Norton, Massachusetts) to send trigger pulses that enabled us to align stimuli onsets and behavioral responses to neural recordings.

A schematic of the task is shown in **Figure 1**. Participants were presented two words, 875 ms presentation time, with a 400 ms blank screen between them. At the end of each trial, participants were asked to indicate via a button press whether the two words were same or different. Word presentation was either visual or auditory. On average, we presented 1500 ± 710 trials (**Table S1** shows the number of trials per participant).

There were four types of trials: Noun followed by Adjective (42% of trials, e.g., “apple green”), Adjective followed by Noun (42% of trials, e.g., “green apple”), Repeated Noun (8% of trials, e.g., “apple apple”), and Repeated Adjective (8% of trials, e.g., “green green”). The order of trials (stimulus presentation modality and noun/adjective structure) was randomly interleaved. Each word combination was presented in a randomized manner 5 times in the audio modality and 5 times in the visual modality. The nouns belonged to two categories, animals (e.g., “cat”) and food (e.g., “apple”). The adjectives belonged to two categories, concrete adjectives (e.g., “big”) and abstract adjectives (e.g., “good”). A list of all the nouns and adjectives is included in **Table S3**. We selected only high frequency English words that were more frequent than 10^-6^ in Google Ngram and were 3 to 7 letters long and had no more than 1 or 2 syllables. We used the max frequency of a word between 2006 and 2019. Finally, we created a balanced selection of nouns and adjectives such that noun and adjectives were indistinguishable from each other using word length or number of syllables (p>0.05 ranksum test). We conducted the experiment in 3 languages, English (16 monolingual and 1 bilingual participant), Spanish (1 bilingual participant) and Taiwanese (3 monolingual participants). Two bilingual international scholars whose native language was Spanish (MAG), and Taiwanese (YLK) translated the words in the task. For non-English languages, we also kept nouns and adjectives indistinguishable based on word-length and number of syllables. Within each language in our data, audio duration was significantly correlated with text length (R² = 0.19–0.25, p < 0.001, **Figure 1h-j**). Word-type comparisons showed no differences in audio duration for any language (t-tests, p>0.05). Only Taiwanese nouns and adjectives differed in text length (t-test, p<0.01), but this did not impact any of the results (see control analyses using General Linear Model (GLM)).

Participants had to indicate whether the two words in a trial were the same or not. The motor responses were the same for nouns or adjectives. The motor responses were also the same for noun followed by adjective or adjective followed by noun trials. Thus, the motor responses were orthogonal to parts of speech and grammar and differences between nouns and adjectives cannot be attributed to motor signals.

### Method Details

#### Preprocessing

A total of 2,428 electrode contacts were implanted, 627 of which were excluded from analysis due to bipolar referencing, presence of line noise or recording artifacts^65^. We removed 60 Hz line noise and its harmonics using a fifth-order Butterworth filter. We focus on the high-gamma band of the intracranial field potential signals obtained by bandpass filtering raw data of each electrode in the 65–150 Hz range (fifth-order Butterworth filter). The high gamma band (65-150 Hz) power was computed using the Chronux toolbox^69^. We used a time-bandwidth product of 3 and 4 leading tapers, a moving window size of 200 ms, and a step size of 5 ms. For every trial, we computed the normalized high gamma activity by subtracting the mean activity from −150 to 50 ms from the onset of the first fixation and then dividing by the standard deviation. This normalized response is reported as “gamma power” on the y-axis when showing electrode responses and referred to throughout the manuscript as *neural responses*.

#### Responsive Electrodes

We evaluated whether an electrode was responsive to visual or auditory stimuli by comparing the response magnitude during the 100 to 400 ms post stimulus onset time to the −400 to −100 ms before stimulus onset (e.g., **Figure S1j-m**). The responsiveness threshold was set using Cohen’s d prime coefficient and based on the number of trials for a statistical power of 80% and p<0.01 (one-tailed z-test). We also computed the time at which the neural signals reach half of the maximum amplitude.

#### Part-of-speech selectivity

We compared the neural responses to nouns versus adjectives. We normalized the responses in each trial by subtracting the mean and dividing by the standard deviation of the high gamma power in the baseline period 100 to 400 ms pre-word1 onset. Periods of significant selective activation were tested using a one-tailed t-test with p<0.05 at each time point to differentiate between nouns and adjectives and were corrected for multiple comparisons with a Benjamini-Hochberg false detection rate (FDR) corrected threshold of q<0.05, separately for auditory and visual trials. After fixing the FDR with q<0.05, an electrode was considered to be selective for part of speech if there was a significant difference between nouns and adjectives for a minimum contiguous window of 65 ms.

#### General Linear Model (GLM)

We created a GLM to tease out the experiment variables that significantly contribute to explaining the responses of a given electrode.

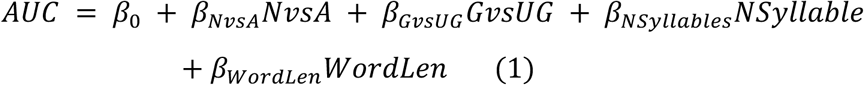

where AUC is the area under the response curve (e.g., **Figure 2a**) from 200 ms to 800 ms after word onset, &_0_is a constant additive term, NvsA is 1 for Nouns and −1 for Adjectives, GvsUG is 1 for Grammatical trials and −1 for Ungrammatical trials, NumberOfSyllables is 1 or 2 (and 0 for visual trials), or WordLength goes from 3 to 7 (and 0 for auditory trials) as the task predictors. We fit this GLM model for each electrode separately using the MATLAB function fitglm and report the corresponding β coefficients (e.g., **Figure 2k**). We assessed whether each coefficient was significantly different from zero when compared to β coefficients generated from shuffled labels (p<0.01, corrected for multiple comparisons).

#### Surprisal Calculation

Surprisal was computed using pre-trained GPT-2 autoregressive language models on Google Colab. The sequence began with a beginning-of-sequence token (BOS), so each pair was encoded as [BOS, w1, w2] using the model’s sub-word tokenizer. Token surprisal was defined as −log p(token | preceding tokens). Word surprisal was summed across tokens for a given word. In a separate analysis, we also added surprisal for each word as a predictor in the GLM.

#### Anatomical comparisons

To assess the degree of anatomical specificity in the neural responses, we compared the percentage of significant electrodes in each brain region to the null distribution expected given the number of electrodes in each area using a permutation test (p<0.01, 10^6^ iterations). A similar approach was followed to compare the same region between the left and right hemispheres.

#### Decoding Analysis

We performed a machine learning decoding analysis to decode parts of speech in individual words combining all the electrodes in each brain region as defined by the Desikan-Killiany atlas^45^ (**Figure 1b-g**). The top-N principal components of all electrodes that explained more than 70% of the variance in the training data for the area under curve of non-overlapping 100 ms time-windows of the signal following word onset were used for decoding. The signal for decoding comprised of features from different frequency bands (beta:12-30 Hz, low gamma:30-65 Hz, and high gamma power: 65-150Hz). The analysis was repeated for 30 random splits of the data with 80% of the data used for training a Support Vector Machine with a linear kernel. Significant decoding performance was found by comparing performance from the original data at each time-window with a null distribution obtained by shuffling labels (p<0.01, ranksum test). Regions with statistically significant decoding performance were found by comparing the average of the maximum decoding performance across time for 30 random iterations of the original data with that of the null distribution, separately for both hemispheres (p<0.01, ranksum test corrected for multiple comparisons) (**Figure 3c,f,i, Figure 4c,f,i,l,o, Figure 5c,f,i**). We also applied a threshold such that for a given region R:

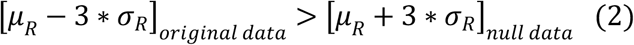

where μ and α represent the average and standard deviation in region R. For the significant regions, the average max-performance between the left and right hemispheres was compared to find if decoding performance was lateralized (p<0.01, ranksum test, corrected for multiple comparisons).

### Quantification and Statistical Analysis

#### Statistical criteria for electrode responsiveness

Electrodes were labeled responsive if high-gamma activity from 100–400 ms post-stimulus exceeded baseline (−400 to −100 ms) using Cohen’s d–based thresholds corresponding to 80% power and p < 0.01 (one-tailed z-test).

#### t-tests for noun vs adjective selectivity and false discovery rate correction

Responses to nouns versus adjectives were compared at each time point using one-tailed t-tests (p < 0.05) separately for auditory and visual trials. Significant time points were corrected using Benjamini–Hochberg FDR (q < 0.05). Selectivity required ≥65 ms of contiguous significance at both words.

#### Ranksum tests

Non-parametric comparisons, including decoding-based regional comparisons and laterality assessments, used Wilcoxon rank-sum tests with a significance threshold of p < 0.01 (multiple-comparison corrected).

#### GLM coefficient significance testing

For each electrode, GLM coefficients were compared to coefficients derived from shuffled-label models. Coefficients exceeding the shuffled distribution at p < 0.01 (corrected for multiple comparisons, ranksum test) were considered significant.

#### Permutation tests for anatomical specificity

The fraction of selective electrodes in each anatomical region was compared to a null distribution obtained by randomly permuting region assignments (10⁶ iterations, p < 0.01, ranksum test, corrected for multiple comparisons).

#### Decoding significance tests

Decoding accuracy in each 100-ms window was compared against a null distribution generated by label shuffling. Time windows and regions were significant when decoding exceeded the null at p < 0.01 (corrected for multiple comparisons, ranksum test)

#### Correlation p-values

Significance of correlations was computed using the standard analytic test in which the correlation coefficient is converted to a t-statistic, and a two-tailed p-value is obtained from the corresponding t-distribution given the sample size.

---

**Fig. S1.**
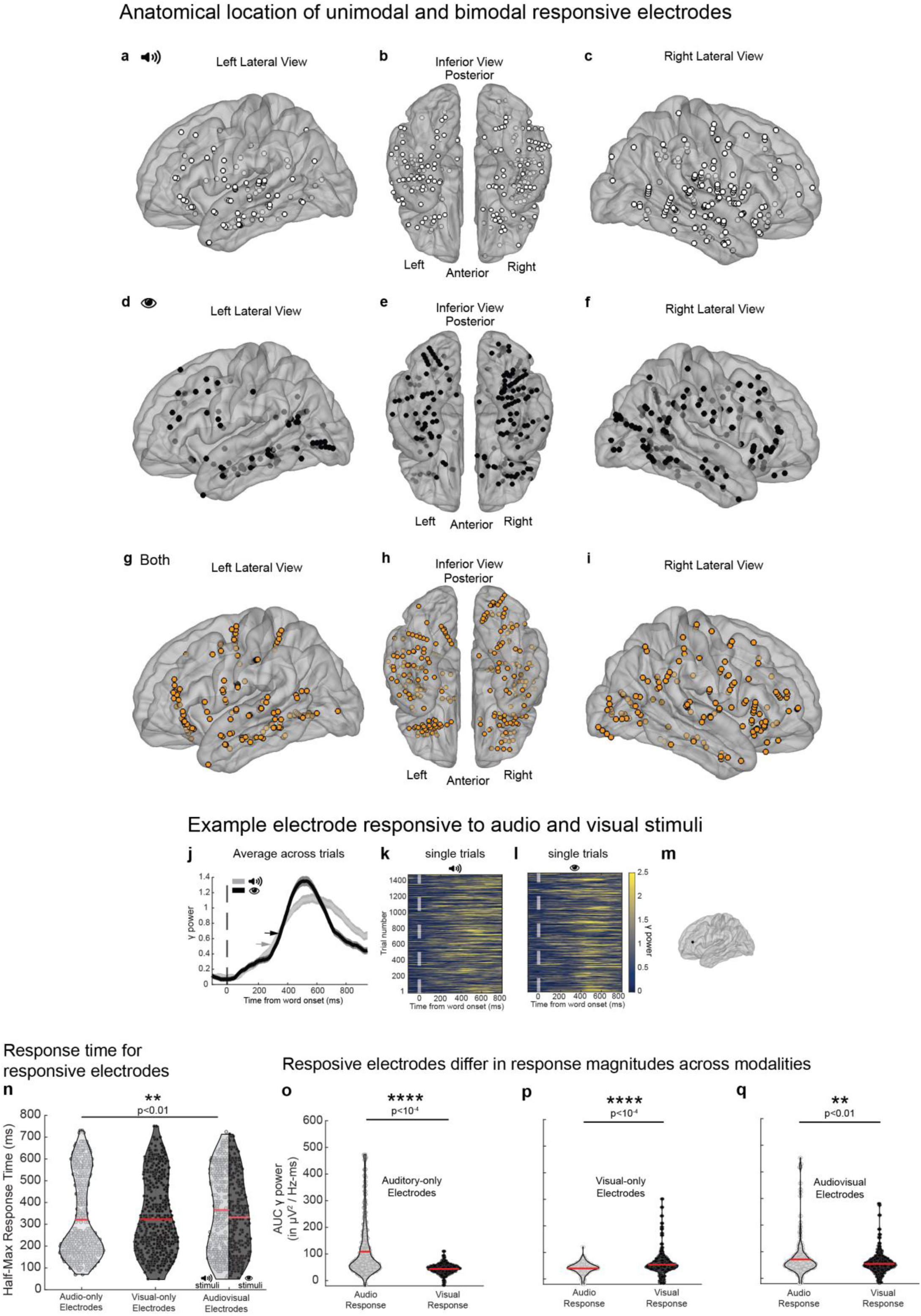
Location of responsive electrodes. **a-c.** Only audio responsive electrodes (**a**: left hemisphere lateral view, n= 102; **b:** inferior view (n=272); **c**: right hemisphere lateral view, n= 170). **d-f.** Only visually responsive electrodes (**d**: n= 85; **e**: n=239; **f:** n= 154). **g-i.** Audiovisual responsive electrodes (**g**: n= 147; **h**: n=293; **i**: n= 146). The same color scheme is followed throughout the paper to indicate vision-only, audio-only or audiovisual electrodes. iELVis pullout factor=20, opaqueness=0.6. **Example responsive electrode. j.** Trial-averaged (± SEM) gamma power for responses to auditory (light grey) or visual (black) presentations for an example electrode in the left rostral middle frontal gyrus (electrode location shown in **m**). Responses are aligned to word onset (vertical dashed line). The arrows indicate the half-maximum time. **k,l**. Raster plots showing each individual trial for the same electrode for each of the 1,496 words for auditor (**k**) and visual (**l**) presentations (see color scale on right). **Half-maximum time and area under the curve for responsive electrodes. n.** Half-maximum time for audio-only electrodes (left, light-gray: 329±187 ms), visual-only electrodes (middle, black: 336±174 ms), and audiovisual electrodes (right; auditory stimuli in light-gray: 379±193 ms, visual stimuli in black: 341±174 ms). There was a small but significant difference between the half-maximum time for auditory-only electrodes and for auditory responses of audiovisual electrodes (p<0.01, ranksum test). Horizontal red bars indicate mean. Horizontal black bars indicate significant differences. **o-q.** Area under the curve for the trial averaged response to auditory stimuli (light-gray violin plots) and visual stimuli (black violin plots) for audio-only electrodes (**o**, auditory stimuli: 108±100 μV^2^/Hz-ms, visual stimuli: 44±16 μV^2^/Hz-ms; p<10^-4^, ranksum test), visual-only electrodes (**p,** auditory stimuli: 40±23 μV^2^/Hz-ms, visual stimuli: 53±43 μV^2^/Hz-ms; p<10^-4^, ranksum test), and audiovisual electrodes (**q,** auditory stimuli: 71±72 μV^2^/Hz-ms, visual stimuli: 54±39 μV^2^/Hz-ms; p<0.01, ranksum test). Horizontal red bars indicate mean. Horizontal black bars indicate significant differences.

**Fig. S2.**
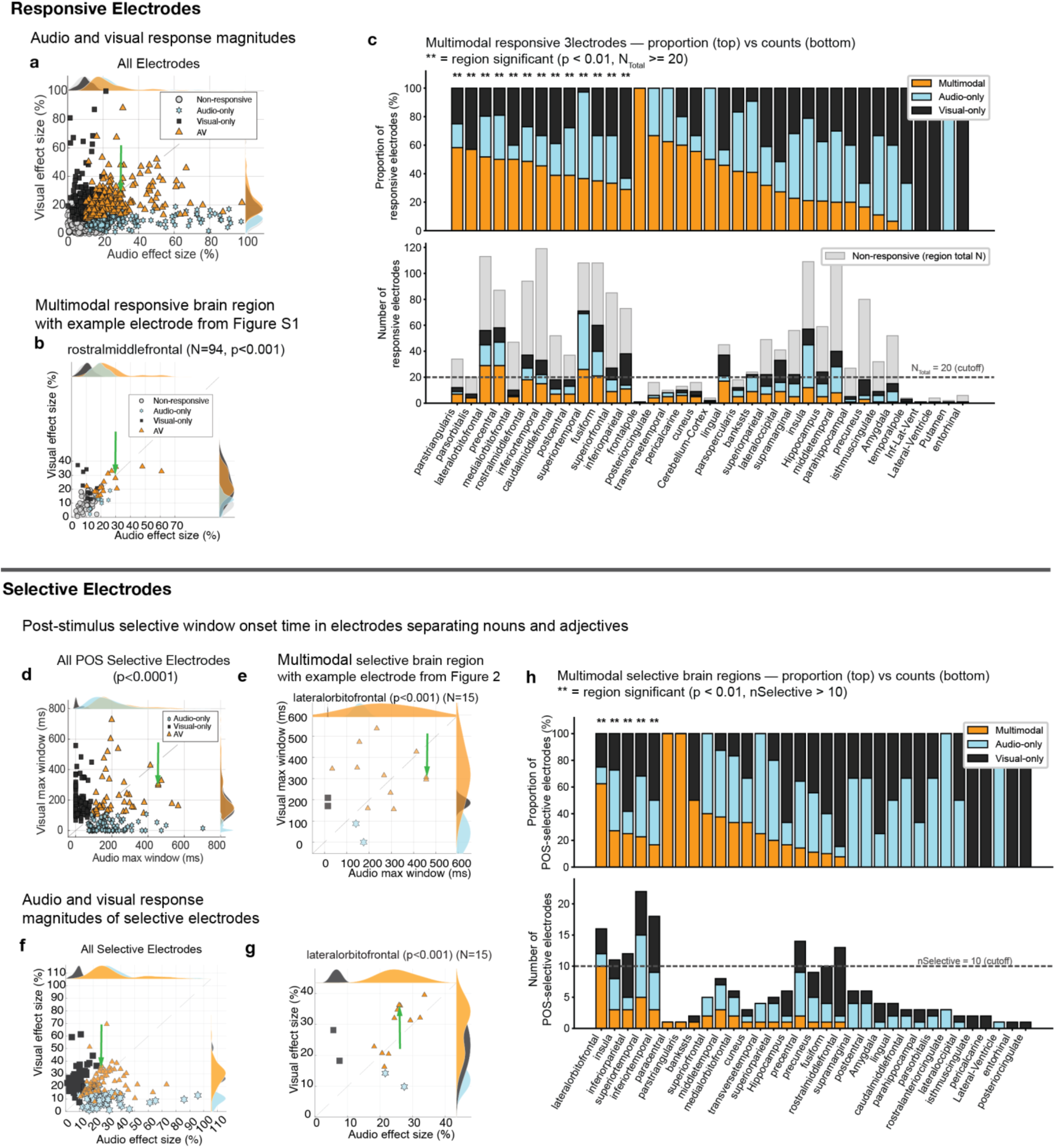
Multimodal responsive regions and parts-of-speech selective regions. **a-n**. **Responsive Electrodes. a.** Scatter plot of all electrodes showing response effect size for auditory trials (x-axis) and visual trials (y-axis). Each electrode can be i) unresponsive (gray circles), ii) auditory-only (cyan hexagram), iii) visual-only (black squares), or iv) audiovisual (orange triangles) responsive (**Methods**). Kernel histograms on either axes show the distribution of responses for each type. **b.** Example region with a significant number of multimodal electrodes (p<0.01, permutation test, n=10^6^ iterations). Green arrow shows location of example electrode in **Figure S1 j-m**. **c.** Brain regions with a significant proportion of multimodal-responsive electrodes, showing the proportion of electrode types (top) and their absolute counts (bottom). Only regions with at least 20 electrodes in total were included (non-responsive in gray). **d-h. Selective Electrodes. d.** Size of maximum contiguous window in milliseconds for selective electrodes. These electrodes differentiate between nouns and adjectives trials for audio-only (n=89, cyan hexagrams), visual-only (n=97, black squares), and audiovisual (n=48, orange triangles) modalities (**Methods**). **e.** Size of maximum contiguous window in milliseconds for selective electrodes in the lateral orbitofrontal region. This region had a significant number of multimodal selective electrodes than expected from the number of audio and visual electrodes (p<0.01, permutation test, n=10^6^ iterations). (**Methods**). **f,g.** Scatter plot of response effect size for selective electrodes across all regions (**f**) and in the lateral orbitofrontal region (**g**). Green arrow shows location of example electrode in **Figure 2 a-h**. **h.** Brain regions with a significant proportion of multimodal-selective electrodes, showing the proportion of electrode types (top) and their absolute counts (bottom). Only regions with at least 10 selective electrodes were included.

**Fig. S3.**
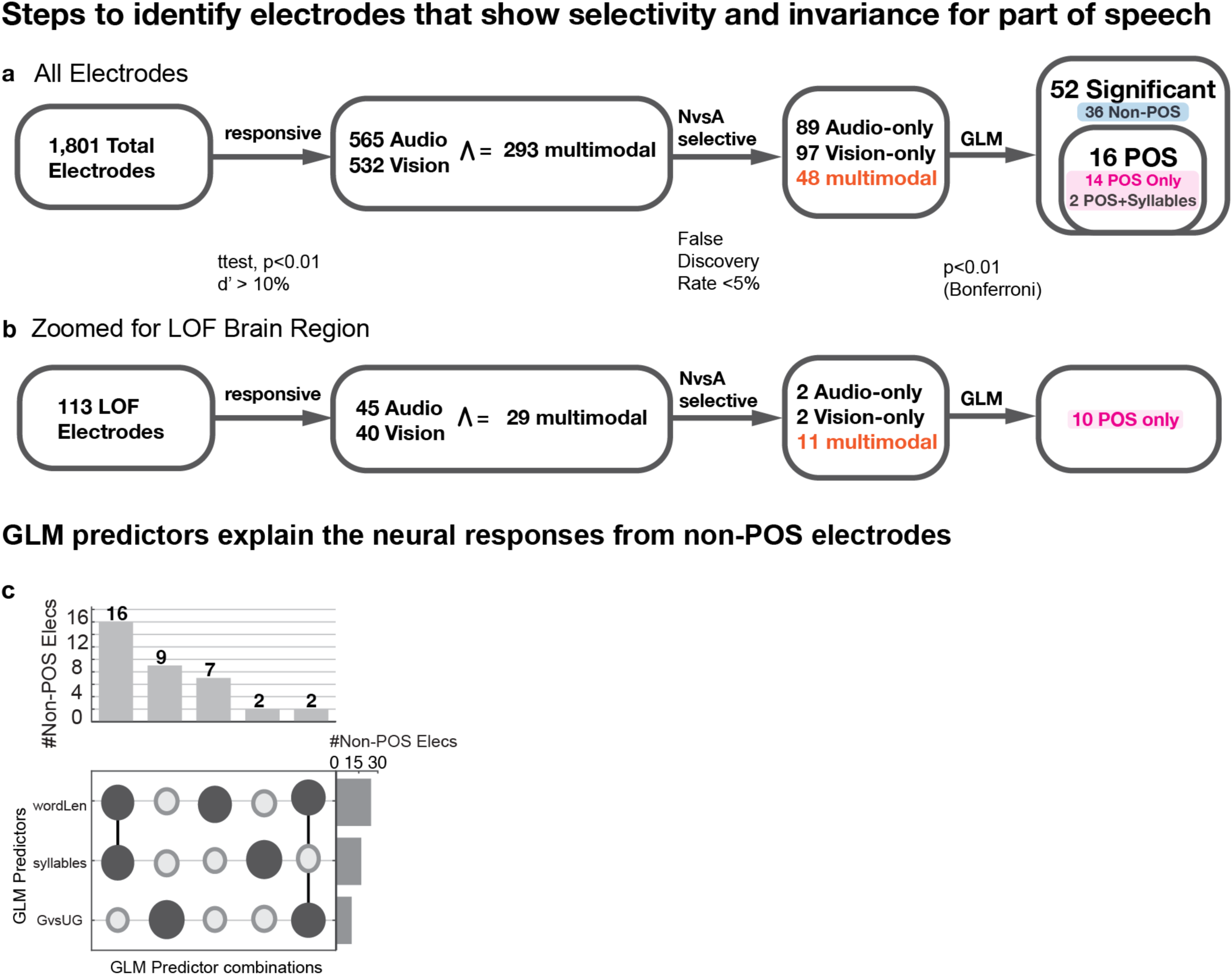
Steps to find parts of speech electrodes. **a,b. Analysis pipeline:** number of electrodes after different stages of analysis: i) total electrodes, ii) responsive electrodes (t-test, p<0.01 and d’ effect size>10%, **Methods**), iii) selective electrodes (t-test, p<0.05, Benjamini-Hochberg false detection rate, q<0.05, **Methods**) and iv) parts-of-speech (p<0.01 only for the NvsA predictor in the GLM, **Methods**) modulated electrodes for the overall dataset (**a**), and for the lateral orbitofrontal region (**b**). A total of 234 (12.99%) were selective for differences between nouns and adjectives for the whole dataset and 15 (13.3% of 113) were selective in the lateral orbitofrontal region (p<0.01 compared to shuffled distribution for both audio and vision, permutation test, n=10^6^ iterations). Finally, a total of 10 electrodes in the lateral orbitofrontal region were significantly predicted by the Nouns versus Adjectives predictor. The 16 POS electrodes in **Figure S2a** include 14 POS-only electrodes and 2 electrodes selective for both POS and syllable count. **c.** Number of significant electrodes predicted by other predictors in the GLM, except Nouns versus Adjectives (Total=36 out of 234; word length= 25; number of syllables=18; Grammaticality=11).

**Fig. S4.**
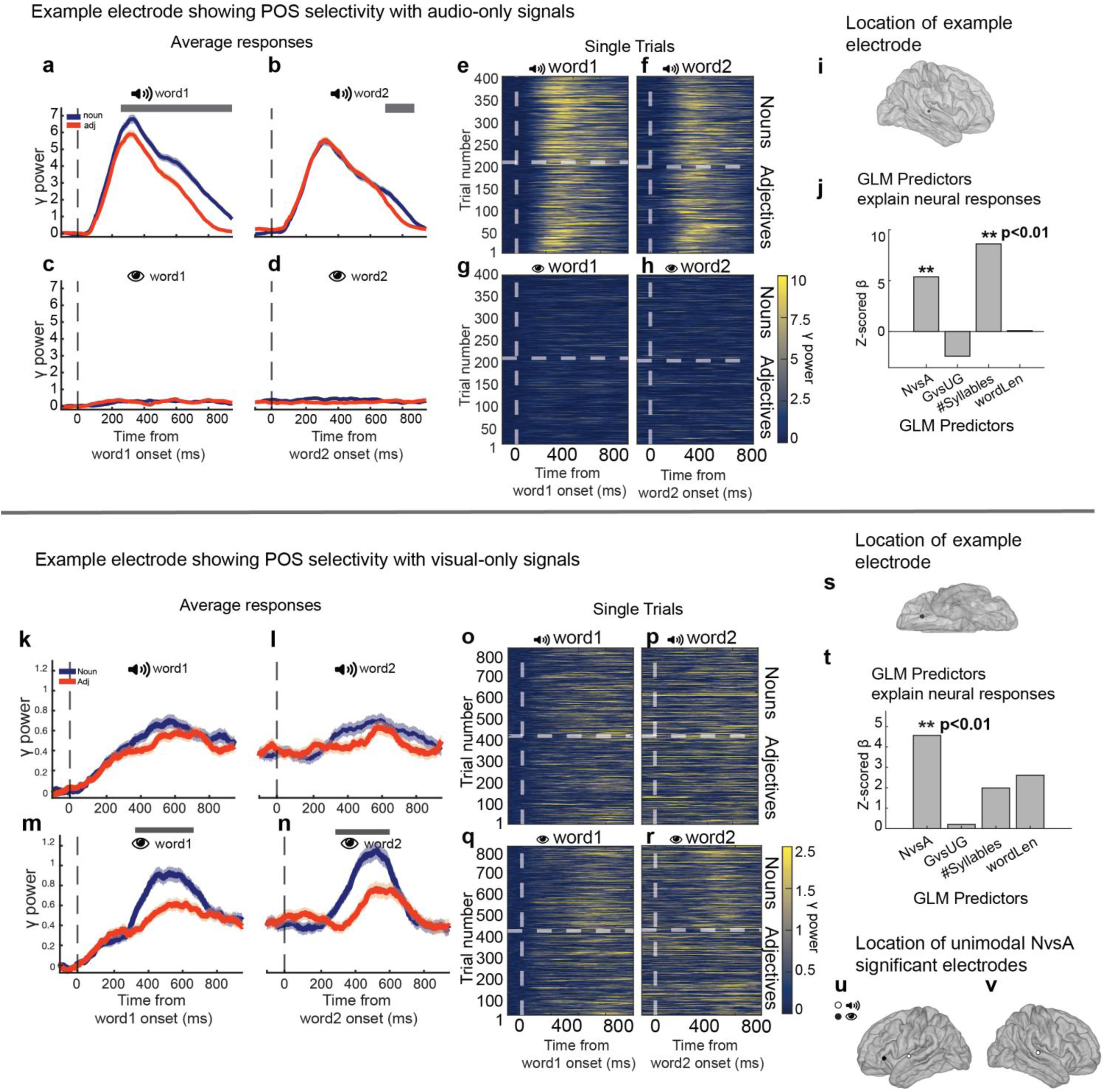
Example electrodes distinguishing parts of speech for only auditory, and only visual stimuli. **a-d.** Trial averaged γ-power of neural responses to Taiwanese words, separated by nouns (blue) and adjectives (red). Neural responses are shown for auditory presentation (**a**, **b**), and visual presentation (**c**, **d**), aligned to word1 onset (**a**, **c**) or word2 onset (**b**, **d**). The vertical dashed lines show word onsets. Shaded areas represent s.e.m. Horizontal lines indicate time periods of statistically significant differences between nouns and adjectives (t-test, p<0.05, Benjamini-Hochberg false detection rate, q<0.05). There was a significant differences between noun and adjectives for auditory presentations shown with a gray horizontal line but no difference for visual presentations. **e-h**. Raster plots showing the responses in individual trials (see color scale on bottom right). **i**. Electrode location in the right insula. **j**. Z-scored β coefficients for Generalized Linear Model used to predict area under the curve between 200 ms and 800 ms post word onset using four task predictors: Noun versus Adjectives, Grammatically Correct versus Ungrammatical, number of syllables (auditory presentation) and word length (visual presentation). Asterisks denote statistically significant coefficients. **k-n.** Trial averaged γ-power of neural responses to Taiwanese words, separated by nouns (blue) and adjectives (red). Neural responses are shown for auditory presentation (**k**, **l**), and visual presentation (**m**, **n**), aligned to word 1 onset (**k**, **m**) or word 2 onset (**l**, **n**). The vertical dashed lines show word onsets. Shaded areas represent s.e.m. Horizontal lines indicate time periods of statistically significant differences between nouns and adjectives (t-test, p<0.05, Benjamini-Hochberg false detection rate, q<0.05). There was a significant differences between noun and adjectives for visual presentations shown with a gray horizontal line but no difference for auditory presentations. **o-r**. Raster plots showing the responses in individual trials (see color scale on bottom right). **s**. Electrode location in the left lateral orbitofrontal. **t**. Z-scored β coefficients for Generalized Linear Model (like **j**). **u,v.** Electrodes in the left **(u)** and right **(v)** hemispheres that showed significant differences between nouns and adjectives either only for auditory trials (white circles) or visual trials (black circles).

**Fig. S5.**
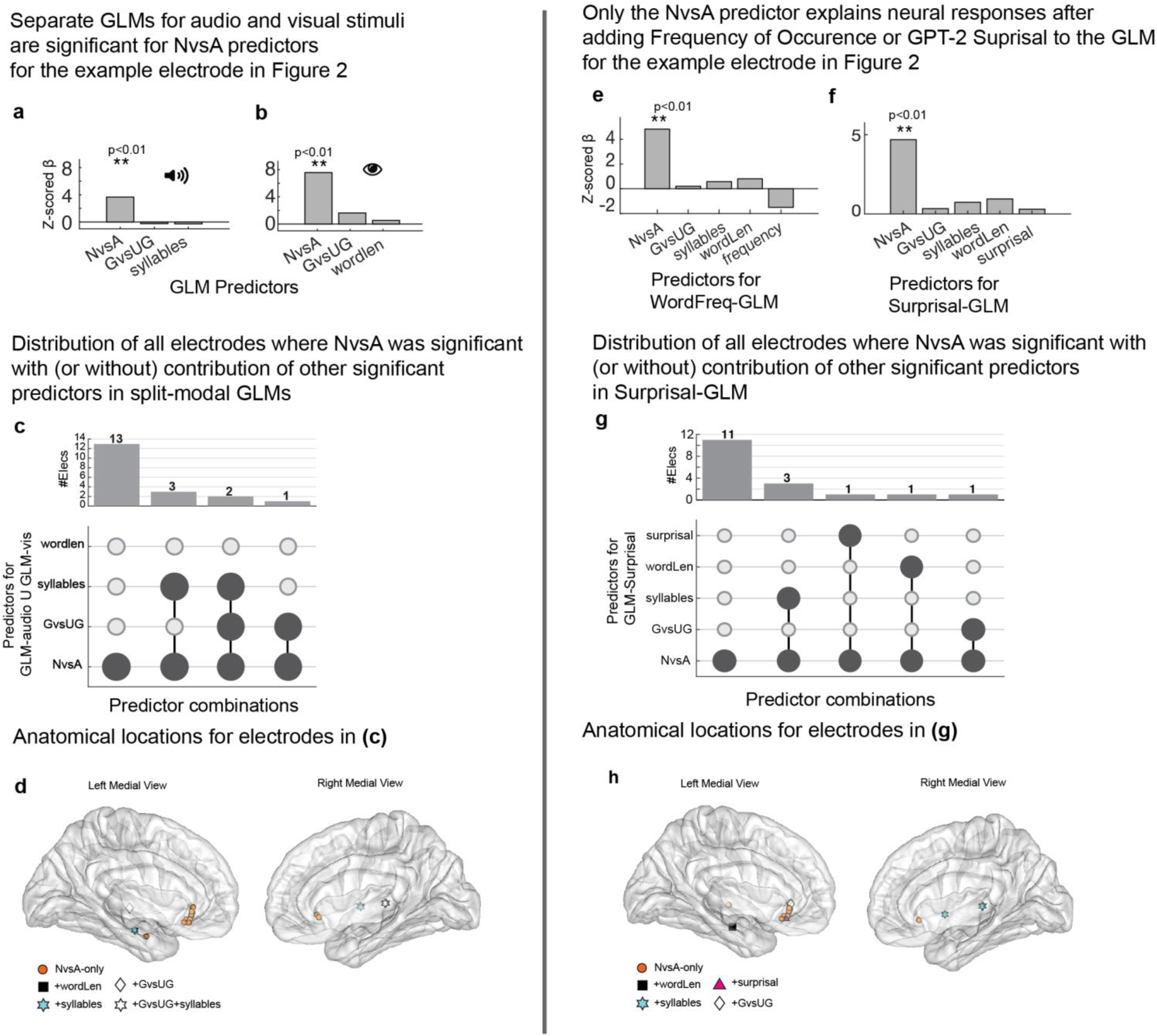
Generalized Linear Models with separate modalities, frequency of occurrence, and surprisal. **a,b.** For the example electrode in **Figure 2,** Z-scored β coefficients for Generalized Linear Model used to predict area under the curve between 200 ms and 800 ms post word onset using three task predictors: Noun versus Adjectives, Grammatically Correct versus Ungrammatical, and number of syllables (**a,** auditory presentation) or word length (**b,** visual presentation). Asterisks denote statistically significant coefficients. **c,d.** POS modulated electrodes that overlapped with other predictors for the modality separated GLMs (shown in **a,b**). Responses from 13 electrodes could only be explained by the NvsA predictor for either audio or visual stimuli (in **d,** orange circles), and 6 other electrodes were also explained by other variables in addition to NvsA (in **d,** n=0, black squares: word length; n=3, cyan hexagrams: number of syllables; n=1, white diamond: grammatically correct or incorrect, n=2, white hexagram: grammatically correct or incorrect and number of syllables). iELVis pullout factor=0, opaqueness=0.4. **e.** For the example electrode in **Figure 2,** Z-scored β coefficients for Generalized Linear Model used to predict area under the curve between 200 ms and 800 ms post word onset with added predictor for frequency of occurrence (natural log of frequency of occurrence was used), compared to **Figure 2j**. **f.** For the example electrode in **Figure 2,** Z-scored β coefficients for Generalized Linear Model used to predict area under the curve between 200 ms and 800 ms post word onset with added predictor for word surprisal, compared to **Figure 2j.** Surprisal values were obtained using probabilities from GPT2 language model. **g,h.** POS modulated electrodes that overlapped with other predictors for the Surprisal-GLM (shown in **e**). Responses from 11 electrodes could only be explained by the NvsA predictor (in **h,** orange circles), and 6 other electrodes were also explained by other variables in addition to NvsA (in **h** n=1, black squares: word length; n=3, cyan hexagrams: number of syllables; n=1, pink triangles: word surprisal; n=1, white diamond: grammatically correct or incorrect). iELVis pullout factor=0, opaqueness=0.4.

**Fig. S6.**
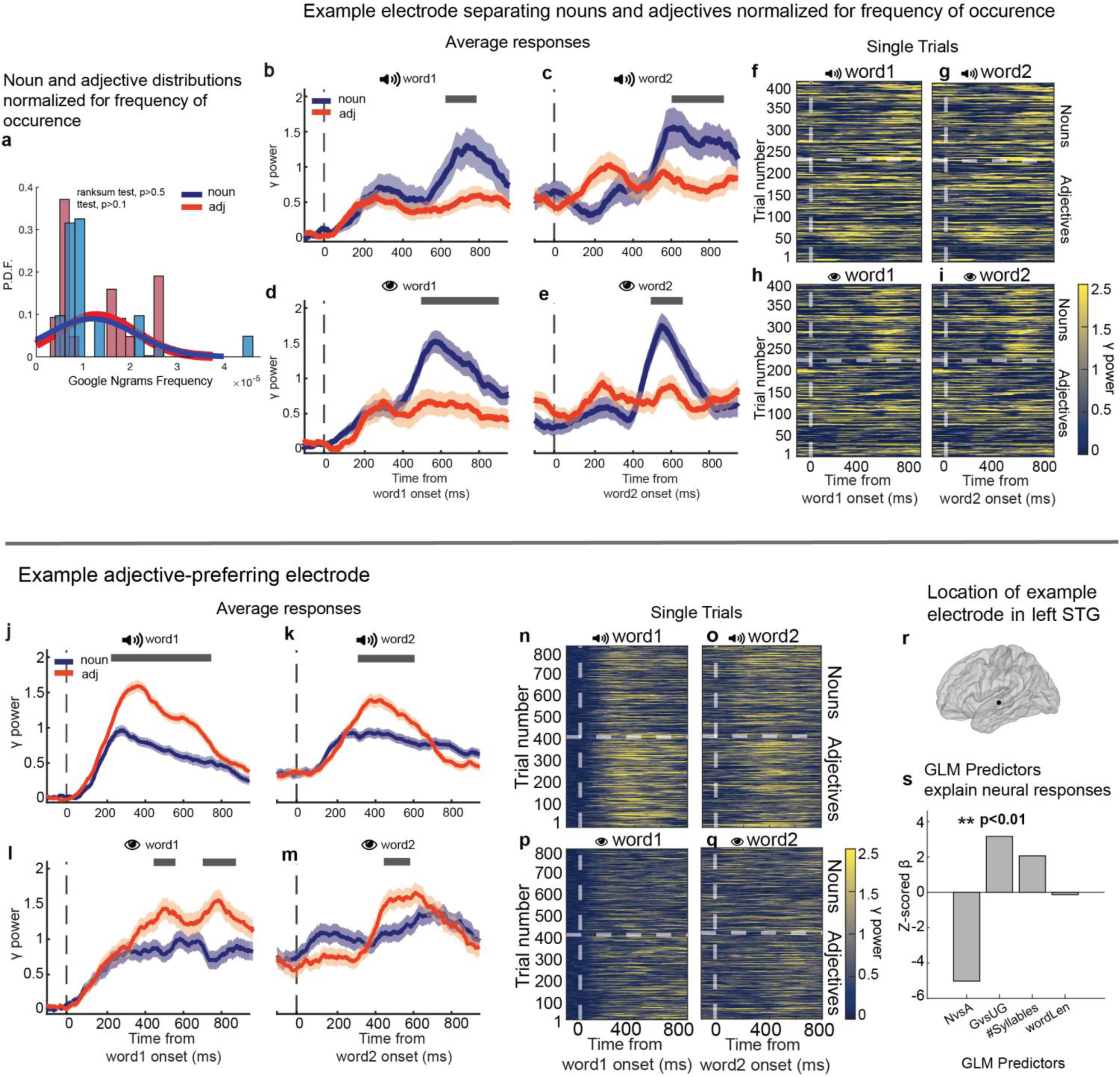
Electrode sensitivity to POS class with frequency control, and another electrode showing adjective preference. Google Ngrams frequency distribution of nouns (blue) and adjectives (red) that were matched for their median (p>0.05, ranksum test) and mean (p>0.05, t-test). **b-e.** Trial averaged γ-power of neural responses to word onsets, separated by nouns (blue) and adjectives (red). Neural responses are shown for auditory presentation (**b**, **c**), and visual presentation (**d**, **e**), aligned to word 1 onset (**b**, **d**) or word 2 onset (**c**, **e**). The vertical dashed lines show word onsets. Shaded areas represent s.e.m. Horizontal lines indicate time periods of statistically significant differences between noun subcategories and adjective subcategories (t-test, p<0.05, Benjamini-Hochberg false detection rate, q<0.05). **f-i**. Raster plots showing the responses in individual trials (see color scale on bottom right). **j-m.** Trial averaged γ-power of neural responses to English words, separated by nouns (blue), and adjectives (red). Neural responses are shown for auditory presentation (**j**, **k,,** n=442 grammatical and 438 ungrammatical trials), and visual presentation (**l**, **m,** n=432 grammatical and 434 ungrammatical trials), aligned to word 1 onset (**j, l**) or word 2 onset (**k, m**). The vertical dashed lines show word onsets. Shaded areas represent s.e.m. Horizontal lines indicate time periods of statistically significant differences between nouns and adjectives (t-test, p<0.05, Benjamini-Hochberg false detection rate, q<0.05). **n-q**. Raster plots showing the responses in individual trials (see color scale on bottom right). **r.** Electrode location in the left superior temporal gyrus. **s**. Z-scored β coefficients for Generalized Linear Model used to predict area under the curve between 200 ms and 800 ms post word using four task predictors: Noun versus Adjectives, Grammatical versus Ungrammatical, number of syllables (auditory presentation) and word length (visual presentation). Asterisks denote statistically significant coefficients. Only the Nouns vs Adjective task predictor was significant and showed a preference for adjectives (p<0.01, corrected for multiple comparisons and β_NvsA_ < 0).

**Fig. S7.**
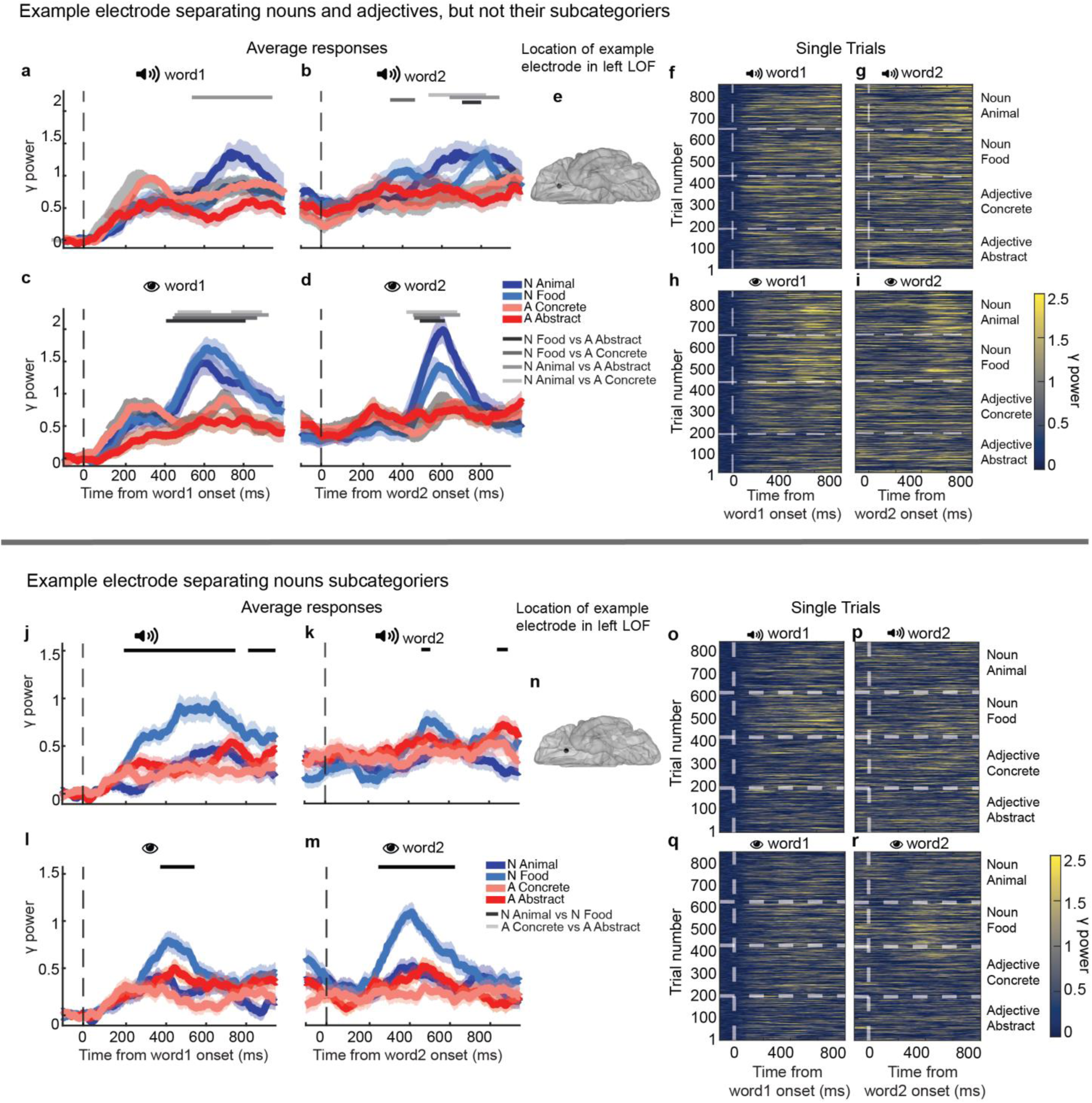
Complementary electrodes showing broad word-class selectivity versus fine-grained noun-subcategory discrimination. **a-d.** Trial averaged γ-power of neural responses to words, separated by animal nouns (dark blue), food nouns (light blue), concrete adjectives (light red), and abstract adjectives (dark red). Neural responses are shown for auditory presentation (**a**, **b**), and visual presentation (**c**, **d**), aligned to word 1 onset (**a**, **c**) or word 2 onset (**b**, **d**). The vertical dashed lines show word onsets. Shaded areas represent s.e.m. Horizontal lines indicate time periods of statistically significant differences between noun subcategories and adjective subcategories (t-test, p<0.05, Benjamini-Hochberg false detection rate, q<0.05). There were no significant differences between noun sub-categories or between adjective sub-categories. **e**. Electrode location in left lateral orbitofrontal. **f-i**. Raster plots showing the responses in individual trials (see color scale on bottom right). **j-m**. Trial averaged γ-power of neural responses to words, separated by animal nouns (dark blue), food nouns (light blue), concrete adjectives (light red), and abstract adjectives (dark red). Neural responses are shown for auditory presentation (**j**, **k**), and visual presentation (**l**, **m**), aligned to word 1 onset (**j**, **l**) or word 2 onset (**k**, **m**). The vertical dashed lines show word onsets. Shaded areas represent s.e.m. Horizontal lines indicate time periods of statistically significant differences between noun subcategories and adjective subcategories (t-test, p<0.05, Benjamini-Hochberg false detection rate, q<0.05). There was a significant differences between noun sub-categories shown with a black horizontal line but no difference between adjective sub-categories. **n.** Electrode location in the left lateral orbitofrontal. **o-r**. Raster plots showing the responses in individual trials (see color scale on bottom right).

**Table S1.** Information about each participant including age, gender, language (ENglish, SPanish, TaiWanese), handedness, number of trials, behavioral performance and number of electrodes.

| Subject | Age | Gender | Language | Handedness | #Trials | %Correct | #Electrodes |
| --- | --- | --- | --- | --- | --- | --- | --- |
| 1 | 9 | M | EN | R | 1178 | 98.5 | 142 |
| 2 | 14 | F | EN | R | 1332 | 94.7 | 100 |
| 3 | 22 | M | EN | R | 1520 | 97.5 | 212 |
| 4 | 49 | F | EN | L | 760 | 83.3 | 44 |
| 5 | 18 | F | EN | R | 3573 | 99.6 | 100 |
| 6 | 20 | M | EN | R | 760 | 99.3 | 139 |
|  |  |  |  |  | 760 | 67.5 |  |
| 7 | 16 | F | EN | L | 760 | 97.9 | 88 |
| 8 | 12 | F | EN | R | 760 | 92.9 | 131 |
| 9 | 37 | F | EN | R | 1900 | 89.3 | 51 |
| 10 | 47 | F | EN | R | 1895 | 98.3 | 75 |
| 11 | 12 | M | EN | R | 950 | 95.5 | 135 |
| 12 | 13 | F | EN | R | 950 | 99.3 | 121 |
| 13 | 25 | M | EN | L write, R throw | 1520 | 99.3 | 84 |
| 14 | 26 | F | EN | L write, R other | 1900 | 97.4 | 32 |
| 15 | 32 | F | EN | R | 1521 | 98.4 | 29 |
|  |  |  |  |  | EN:950 |  |  |
| 16 | 22 | M | EN & SP | R | SP:950 | 97.3 | 77 |
| 17 | 42 | M | TW | R | 1068 | 89 | 57 |
| 18 | 36 | M | TW | R | 2429 | 87.1 | 59 |
| 19 | 53 | F | EN | R | 950 | 90.5 | 73 |
| 20 | 44 | M | TW | R | 1900 | NA | 52 |
| TOTAL |  |  |  |  |  |  | 1801 |

**Table S2.** Distribution of electrodes over the Desikan-Killiany Atlas. The number of electrodes for different brain regions of the DK atlas (rows) for different conditions (columns). From the left to right the columns represent the following: (1) Total electrodes, (2) Gray Matter Left, (3) White Matter Left, (4) Total Left, (5) Gray Matter Right, (6) White Matter Right, (7) Total Right, (8) Responsive Audio, (9) Responsive Visual, (10) Responsive Audiovisual. The regions that showed a significant percent of audiovisual electrodes that was statistically unlikely to get from a random intersection of audio or visual electrodes are highlighted in bold (p<0.01, permutation test, n=10^6^ iterations, total electrodes >=20)

| Region\nelecs | Total | GMLeft | WMLeft | Left | GMRight | WMRight | Right | rAud | rVis | rAV |
| --- | --- | --- | --- | --- | --- | --- | --- | --- | --- | --- |
| Amygdala | 52 | 21 | 0 | 21 | 31 | 0 | 31 | 9 | 7 | 1 |
| Cerebellum-Cortex | 4 | 3 | 0 | 3 | 1 | 0 | 1 | 2 | 1 | 1 |
| Hippocampus | 59 | 33 | 0 | 33 | 26 | 0 | 26 | 15 | 14 | 5 |
| Inf-Lat-Vent | 1 | 1 | 0 | 1 | 0 | 0 | 0 | 0 | 1 | 0 |
| Lateral-Ventricle | 4 | 0 | 0 | 0 | 4 | 0 | 4 | 0 | 1 | 0 |
| Putamen | 2 | 1 | 0 | 1 | 1 | 0 | 1 | 1 | 0 | 0 |
| VentralDC | 1 | 1 | 0 | 1 | 0 | 0 | 0 | 0 | 0 | 0 |
| bankssts | 24 | 2 | 4 | 6 | 9 | 9 | 18 | 20 | 11 | 9 |
| caudalanteriorcingulate | 17 | 3 | 1 | 4 | 3 | 10 | 13 | 0 | 0 | 0 |
| **caudalmiddlefrontal** | 52 | 2 | 12 | 14 | 21 | 17 | 38 | 11 | 14 | 7 |
| cuneus | 16 | 0 | 3 | 3 | 10 | 3 | 13 | 6 | 8 | 5 |
| entorhinal | 6 | 1 | 1 | 2 | 2 | 2 | 4 | 0 | 1 | 0 |
| frontalpole | 1 | 0 | 0 | 0 | 1 | 0 | 1 | 1 | 1 | 1 |
| **fusiform** | 108 | 19 | 32 | 51 | 27 | 30 | 57 | 40 | 41 | 21 |
| **inferiorparietal** | 73 | 3 | 17 | 20 | 31 | 22 | 53 | 14 | 35 | 11 |
| **inferiortemporal** | 119 | 14 | 38 | 52 | 31 | 36 | 67 | 22 | 26 | 15 |
| insula | 109 | 18 | 29 | 47 | 29 | 33 | 62 | 45 | 24 | 12 |
| isthmuscingulate | 32 | 7 | 4 | 11 | 11 | 10 | 21 | 6 | 4 | 1 |
| lateraloccipital | 41 | 2 | 8 | 10 | 13 | 18 | 31 | 16 | 26 | 9 |
| **lateralorbitofrontal** | 113 | 7 | 31 | 38 | 38 | 37 | 75 | 45 | 40 | 29 |
| lingual | 45 | 6 | 13 | 19 | 8 | 18 | 26 | 21 | 33 | 17 |
| **medialorbitofrontal** | 47 | 10 | 8 | 18 | 14 | 15 | 29 | 6 | 9 | 5 |
| middletemporal | 117 | 27 | 39 | 66 | 24 | 27 | 51 | 28 | 20 | 8 |
| paracentral | 4 | 0 | 0 | 0 | 2 | 2 | 4 | 0 | 0 | 0 |
| parahippocampal | 27 | 12 | 4 | 16 | 6 | 5 | 11 | 3 | 3 | 1 |
| parsopercularis | 18 | 5 | 4 | 9 | 4 | 5 | 9 | 10 | 7 | 5 |
| **parsorbitalis** | 20 | 2 | 5 | 7 | 7 | 6 | 13 | 4 | 7 | 4 |
| **parstriangularis** | 34 | 3 | 4 | 7 | 16 | 11 | 27 | 9 | 10 | 7 |
| pericalcarine | 13 | 3 | 2 | 5 | 3 | 5 | 8 | 8 | 8 | 6 |
| **postcentral** | 37 | 2 | 6 | 8 | 10 | 19 | 29 | 13 | 12 | 7 |
| posteriorcingulate | 16 | 4 | 3 | 7 | 1 | 8 | 9 | 6 | 4 | 4 |
| **precentral** | 87 | 10 | 10 | 20 | 33 | 34 | 67 | 47 | 40 | 29 |
| precuneus | 80 | 12 | 24 | 36 | 14 | 30 | 44 | 6 | 15 | 3 |
| rostralanteriorcingulate | 16 | 3 | 4 | 7 | 5 | 4 | 9 | 0 | 0 | 0 |
| **rostralmiddlefrontal** | 94 | 16 | 22 | 38 | 24 | 32 | 56 | 27 | 28 | 18 |
| **superiorfrontal** | 85 | 17 | 16 | 33 | 19 | 33 | 52 | 18 | 18 | 9 |
| superiorparietal | 49 | 4 | 6 | 10 | 15 | 24 | 39 | 13 | 16 | 7 |
| **superiortemporal** | 108 | 20 | 38 | 58 | 15 | 35 | 50 | 69 | 28 | 26 |
| supramarginal | 56 | 0 | 0 | 0 | 30 | 26 | 56 | 15 | 12 | 5 |
| temporalpole | 4 | 3 | 1 | 4 | 0 | 0 | 0 | 1 | 2 | 0 |
| transverse temporal | 10 | 2 | 5 | 7 | 2 | 1 | 3 | 8 | 5 | 5 |
| **TOTAL** | **1801** | **299** | **394** | **693** | **541** | **567** | **1108** | **565** | **532** | **293** |

**Table S3.**
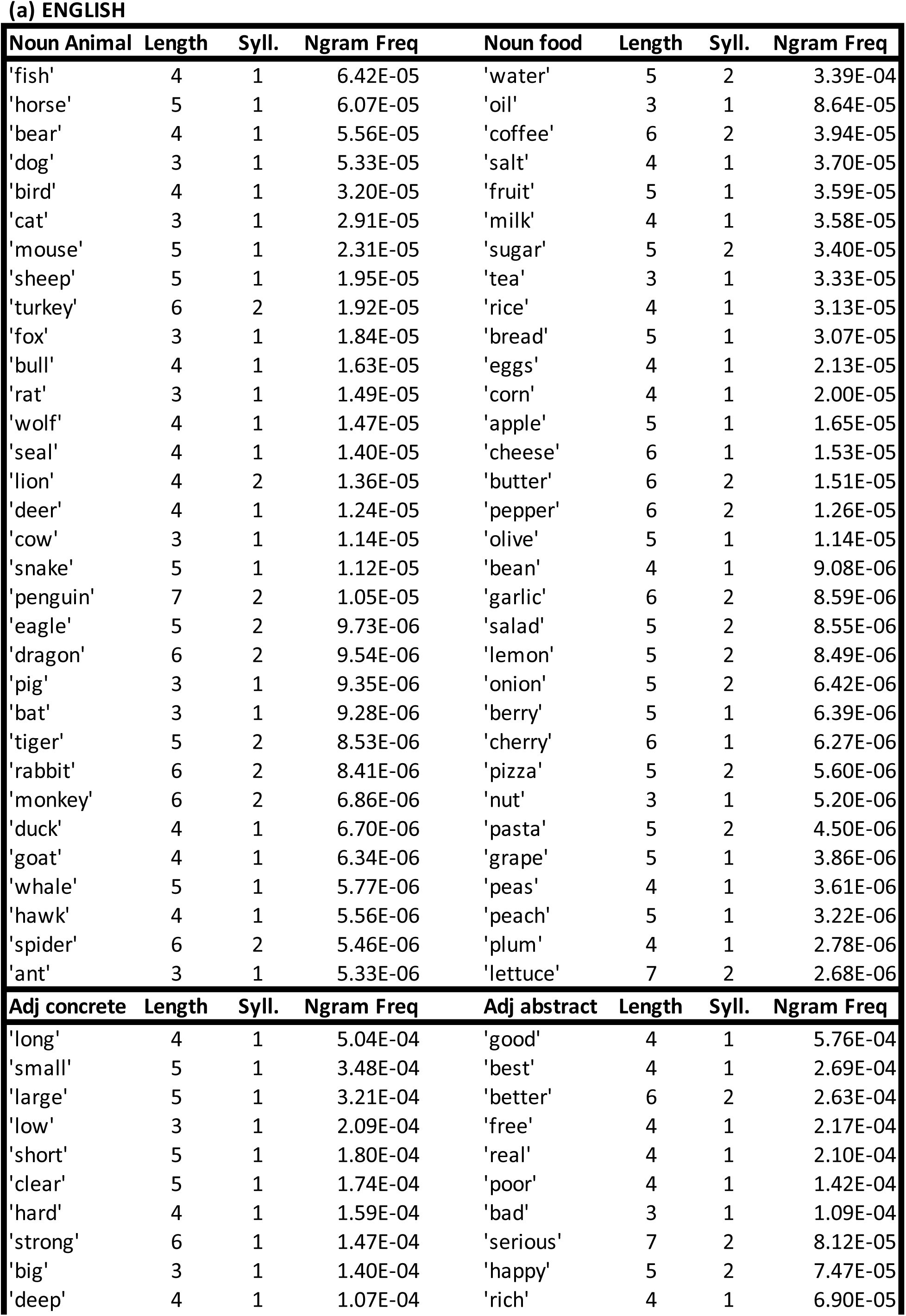
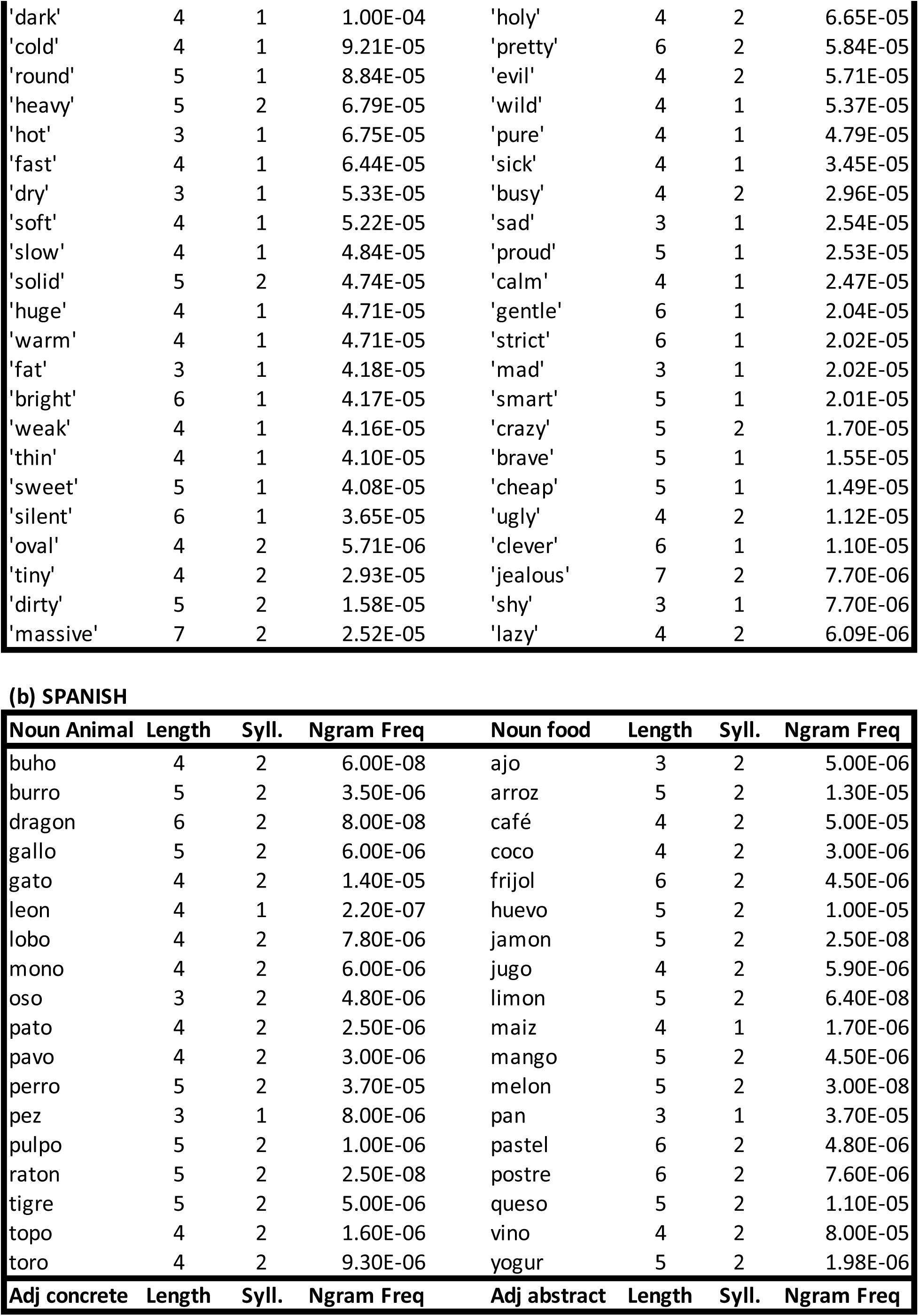
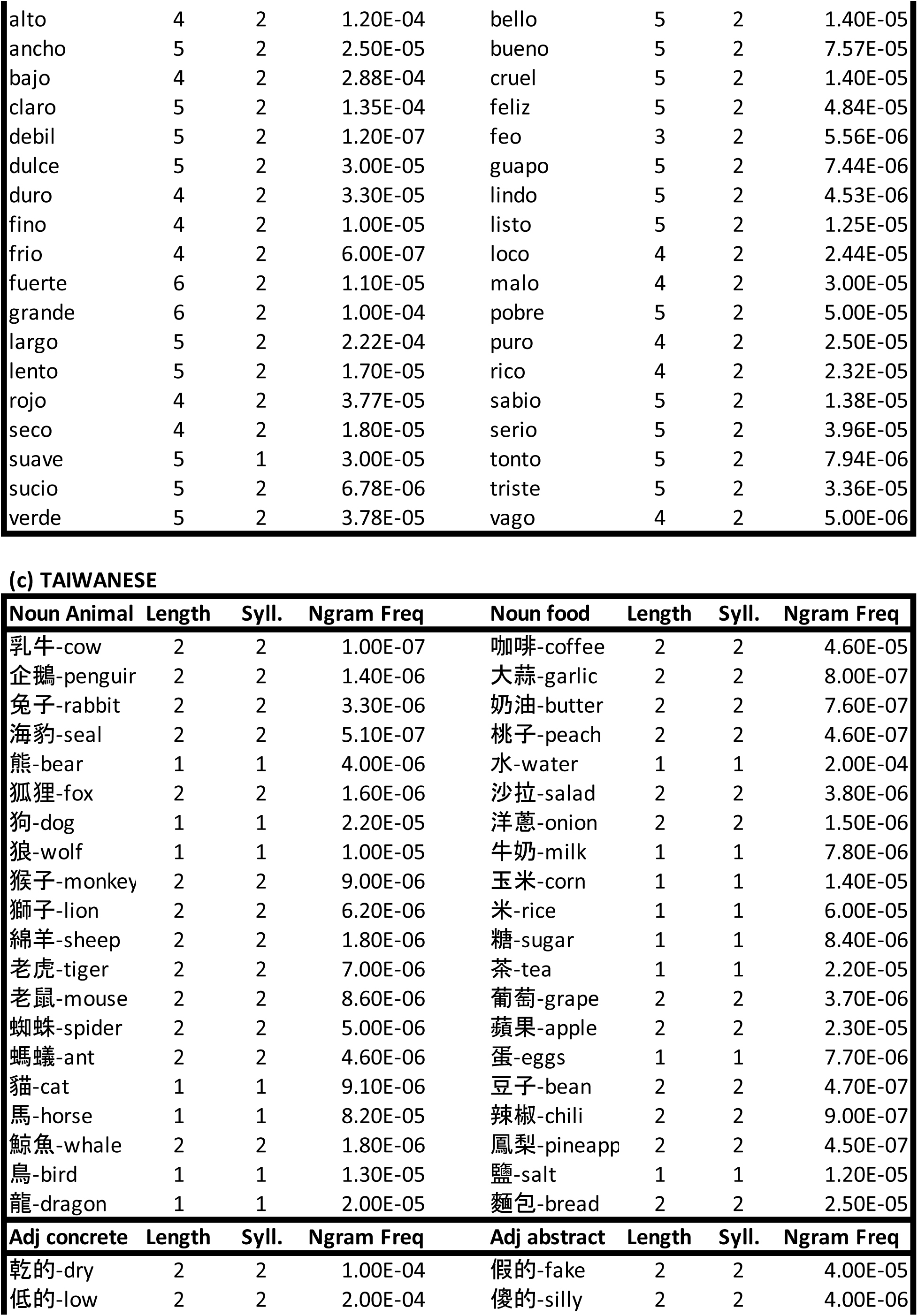
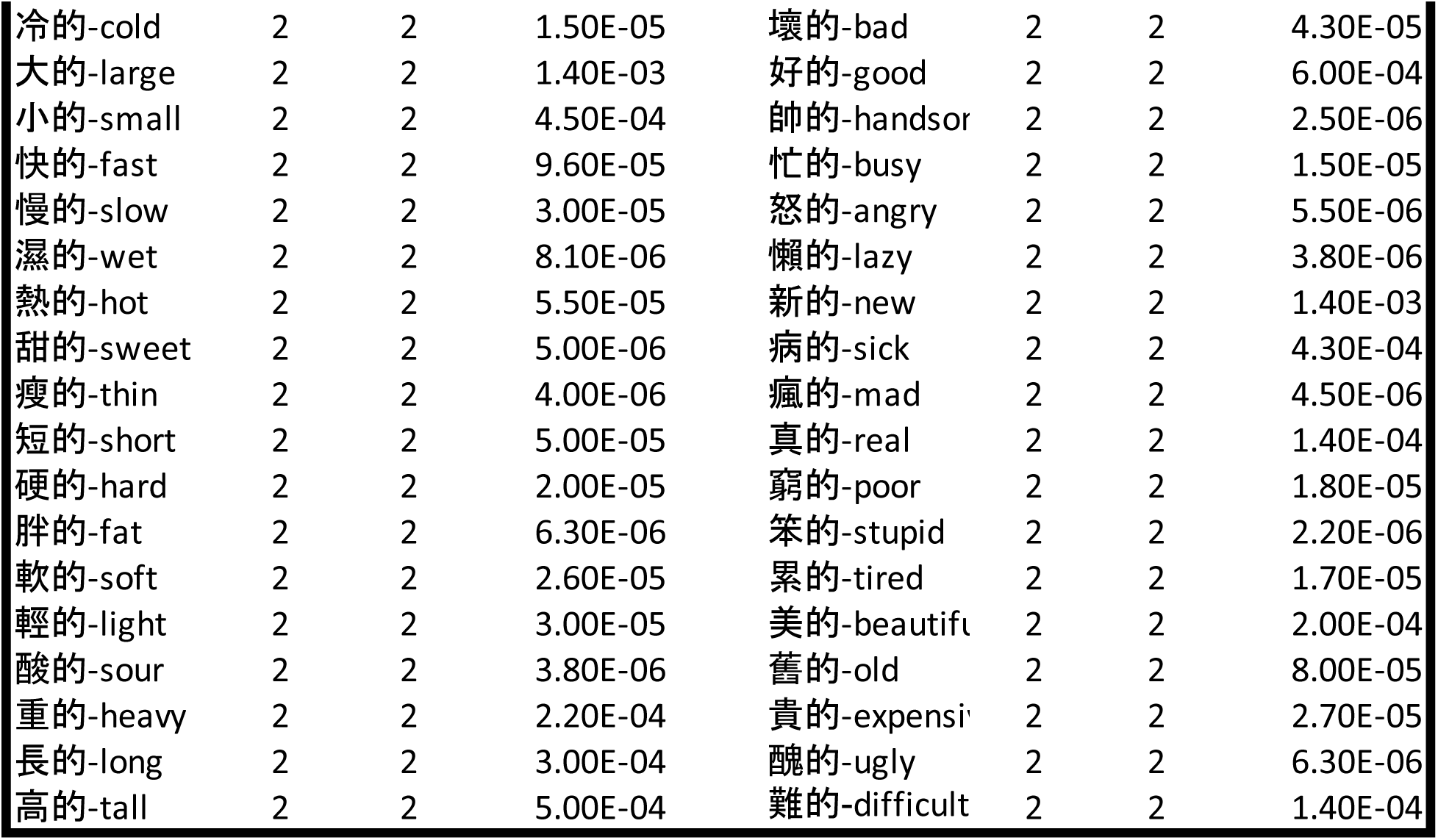
List of all the words used in the experiment, their lengths, number of syllables, and occurrence frequency. (a) English. (b) Spanish. (c) Taiwanese

**(a) ENGLISH**
| **Noun Animal** | **Length** | **Syll.** | **Ngram Freq** | **Noun food** | **Length** | **Syll.** | **Ngram Freq** |
| --- | --- | --- | --- | --- | --- | --- | --- |
| 'fish' | 4 | 1 | 6.42E-05 | 'water' | 5 | 2 | 3.39E-04 |
| 'horse' | 5 | 1 | 6.07E-05 | 'oil' | 3 | 1 | 8.64E-05 |
| 'bear' | 4 | 1 | 5.56E-05 | 'coffee' | 6 | 2 | 3.94E-05 |
| 'dog' | 3 | 1 | 5.33E-05 | 'salt' | 4 | 1 | 3.70E-05 |
| 'bird' | 4 | 1 | 3.20E-05 | 'fruit' | 5 | 1 | 3.59E-05 |
| 'cat' | 3 | 1 | 2.91E-05 | 'milk' | 4 | 1 | 3.58E-05 |
| 'mouse' | 5 | 1 | 2.31E-05 | 'sugar' | 5 | 2 | 3.40E-05 |
| 'sheep' | 5 | 1 | 1.95E-05 | 'tea' | 3 | 1 | 3.33E-05 |
| 'turkey' | 6 | 2 | 1.92E-05 | 'rice' | 4 | 1 | 3.13E-05 |
| 'fox' | 3 | 1 | 1.84E-05 | 'bread' | 5 | 1 | 3.07E-05 |
| 'bull' | 4 | 1 | 1.63E-05 | 'eggs' | 4 | 1 | 2.13E-05 |
| 'rat' | 3 | 1 | 1.49E-05 | 'corn' | 4 | 1 | 2.00E-05 |
| 'wolf' | 4 | 1 | 1.47E-05 | 'apple' | 5 | 1 | 1.65E-05 |
| 'seal' | 4 | 1 | 1.40E-05 | 'cheese' | 6 | 1 | 1.53E-05 |
| 'lion' | 4 | 2 | 1.36E-05 | 'butter' | 6 | 2 | 1.51E-05 |
| 'deer' | 4 | 1 | 1.24E-05 | 'pepper' | 6 | 2 | 1.26E-05 |
| 'cow' | 3 | 1 | 1.14E-05 | 'olive' | 5 | 1 | 1.14E-05 |
| 'snake' | 5 | 1 | 1.12E-05 | 'bean' | 4 | 1 | 9.08E-06 |
| 'penguin' | 7 | 2 | 1.05E-05 | 'garlic' | 6 | 2 | 8.59E-06 |
| 'eagle' | 5 | 2 | 9.73E-06 | 'salad' | 5 | 2 | 8.55E-06 |
| 'dragon' | 6 | 2 | 9.54E-06 | 'lemon' | 5 | 2 | 8.49E-06 |
| 'pig' | 3 | 1 | 9.35E-06 | 'onion' | 5 | 2 | 6.42E-06 |
| 'bat' | 3 | 1 | 9.28E-06 | 'berry' | 5 | 1 | 6.39E-06 |
| 'tiger' | 5 | 2 | 8.53E-06 | 'cherry' | 6 | 1 | 6.27E-06 |
| 'rabbit' | 6 | 2 | 8.41E-06 | 'pizza' | 5 | 2 | 5.60E-06 |
| 'monkey' | 6 | 2 | 6.86E-06 | 'nut' | 3 | 1 | 5.20E-06 |
| 'duck' | 4 | 1 | 6.70E-06 | 'pasta' | 5 | 2 | 4.50E-06 |
| 'goat' | 4 | 1 | 6.34E-06 | 'grape' | 5 | 1 | 3.86E-06 |
| 'whale' | 5 | 1 | 5.77E-06 | 'peas' | 4 | 1 | 3.61E-06 |
| 'hawk' | 4 | 1 | 5.56E-06 | 'peach' | 5 | 1 | 3.22E-06 |
| 'spider' | 6 | 2 | 5.46E-06 | 'plum' | 4 | 1 | 2.78E-06 |
| 'ant' | 3 | 1 | 5.33E-06 | 'lettuce' | 7 | 2 | 2.68E-06 |
| **Adj concrete** | **Length** | **Syll.** | **Ngram Freq** | **Adj abstract** | **Length** | **Syll.** | **Ngram Freq** |
| 'long' | 4 | 1 | 5.04E-04 | 'good' | 4 | 1 | 5.76E-04 |
| 'small' | 5 | 1 | 3.48E-04 | 'best' | 4 | 1 | 2.69E-04 |
| 'large' | 5 | 1 | 3.21E-04 | 'better' | 6 | 2 | 2.63E-04 |
| 'low' | 3 | 1 | 2.09E-04 | 'free' | 4 | 1 | 2.17E-04 |
| 'short' | 5 | 1 | 1.80E-04 | 'real' | 4 | 1 | 2.10E-04 |
| 'clear' | 5 | 1 | 1.74E-04 | 'poor' | 4 | 1 | 1.42E-04 |
| 'hard' | 4 | 1 | 1.59E-04 | 'bad' | 3 | 1 | 1.09E-04 |
| 'strong' | 6 | 1 | 1.47E-04 | 'serious' | 7 | 2 | 8.12E-05 |
| 'big' | 3 | 1 | 1.40E-04 | 'happy' | 5 | 2 | 7.47E-05 |
| 'deep' | 4 | 1 | 1.07E-04 | 'rich' | 4 | 1 | 6.90E-05 |
| 'dark' | 4 | 1 | 1.00E-04 | 'holy' | 4 | 2 | 6.65E-05 |
| 'cold' | 4 | 1 | 9.21E-05 | 'pretty' | 6 | 2 | 5.84E-05 |
| 'round' | 5 | 1 | 8.84E-05 | 'evil' | 4 | 2 | 5.71E-05 |
| 'heavy' | 5 | 2 | 6.79E-05 | 'wild' | 4 | 1 | 5.37E-05 |
| 'hot' | 3 | 1 | 6.75E-05 | 'pure' | 4 | 1 | 4.79E-05 |
| 'fast' | 4 | 1 | 6.44E-05 | 'sick' | 4 | 1 | 3.45E-05 |
| 'dry' | 3 | 1 | 5.33E-05 | 'busy' | 4 | 2 | 2.96E-05 |
| 'soft' | 4 | 1 | 5.22E-05 | 'sad' | 3 | 1 | 2.54E-05 |
| 'slow' | 4 | 1 | 4.84E-05 | 'proud' | 5 | 1 | 2.53E-05 |
| 'solid' | 5 | 2 | 4.74E-05 | 'calm' | 4 | 1 | 2.47E-05 |
| 'huge' | 4 | 1 | 4.71E-05 | 'gentle' | 6 | 1 | 2.04E-05 |
| 'warm' | 4 | 1 | 4.71E-05 | 'strict' | 6 | 1 | 2.02E-05 |
| 'fat' | 3 | 1 | 4.18E-05 | 'mad' | 3 | 1 | 2.02E-05 |
| 'bright' | 6 | 1 | 4.17E-05 | 'smart' | 5 | 1 | 2.01E-05 |
| 'weak' | 4 | 1 | 4.16E-05 | 'crazy' | 5 | 2 | 1.70E-05 |
| 'thin' | 4 | 1 | 4.10E-05 | 'brave' | 5 | 1 | 1.55E-05 |
| 'sweet' | 5 | 1 | 4.08E-05 | 'cheap' | 5 | 1 | 1.49E-05 |
| 'silent' | 6 | 1 | 3.65E-05 | 'ugly' | 4 | 2 | 1.12E-05 |
| 'oval' | 4 | 2 | 5.71E-06 | 'clever' | 6 | 1 | 1.10E-05 |
| 'tiny' | 4 | 2 | 2.93E-05 | 'jealous' | 7 | 2 | 7.70E-06 |
| 'dirty' | 5 | 2 | 1.58E-05 | 'shy' | 3 | 1 | 7.70E-06 |
| 'massive' | 7 | 2 | 2.52E-05 | 'lazy' | 4 | 2 | 6.09E-06 |

**(b) SPANISH**
| **Noun Animal** | **Length** | **Syll.** | **Ngram Freq** | **Noun food** | **Length** | **Syll.** | **Ngram Freq** |
| --- | --- | --- | --- | --- | --- | --- | --- |
| buho | 4 | 2 | 6.00E-08 | ajo | 3 | 2 | 5.00E-06 |
| burro | 5 | 2 | 3.50E-06 | arroz | 5 | 2 | 1.30E-05 |
| dragon | 6 | 2 | 8.00E-08 | café | 4 | 2 | 5.00E-05 |
| gallo | 5 | 2 | 6.00E-06 | coco | 4 | 2 | 3.00E-06 |
| gato | 4 | 2 | 1.40E-05 | frijol | 6 | 2 | 4.50E-06 |
| leon | 4 | 1 | 2.20E-07 | huevo | 5 | 2 | 1.00E-05 |
| lobo | 4 | 2 | 7.80E-06 | jamon | 5 | 2 | 2.50E-08 |
| mono | 4 | 2 | 6.00E-06 | jugo | 4 | 2 | 5.90E-06 |
| oso | 3 | 2 | 4.80E-06 | limon | 5 | 2 | 6.40E-08 |
| pato | 4 | 2 | 2.50E-06 | maiz | 4 | 1 | 1.70E-06 |
| pavo | 4 | 2 | 3.00E-06 | mango | 5 | 2 | 4.50E-06 |
| perro | 5 | 2 | 3.70E-05 | melon | 5 | 2 | 3.00E-08 |
| pez | 3 | 1 | 8.00E-06 | pan | 3 | 1 | 3.70E-05 |
| pulpo | 5 | 2 | 1.00E-06 | pastel | 6 | 2 | 4.80E-06 |
| raton | 5 | 2 | 2.50E-08 | postre | 6 | 2 | 7.60E-06 |
| tigre | 5 | 2 | 5.00E-06 | queso | 5 | 2 | 1.10E-05 |
| topo | 4 | 2 | 1.60E-06 | vino | 4 | 2 | 8.00E-05 |
| toro | 4 | 2 | 9.30E-06 | yogur | 5 | 2 | 1.98E-06 |
| **Adj concrete** | **Length** | **Syll.** | **Ngram Freq** | **Adj abstract** | **Length** | **Syll.** | **Ngram Freq** |
| alto | 4 | 2 | 1.20E-04 | bello | 5 | 2 | 1.40E-05 |
| ancho | 5 | 2 | 2.50E-05 | bueno | 5 | 2 | 7.57E-05 |
| bajo | 4 | 2 | 2.88E-04 | cruel | 5 | 2 | 1.40E-05 |
| claro | 5 | 2 | 1.35E-04 | feliz | 5 | 2 | 4.84E-05 |
| debil | 5 | 2 | 1.20E-07 | feo | 3 | 2 | 5.56E-06 |
| dulce | 5 | 2 | 3.00E-05 | guapo | 5 | 2 | 7.44E-06 |
| duro | 4 | 2 | 3.30E-05 | lindo | 5 | 2 | 4.53E-06 |
| fino | 4 | 2 | 1.00E-05 | listo | 5 | 2 | 1.25E-05 |
| frio | 4 | 2 | 6.00E-07 | loco | 4 | 2 | 2.44E-05 |
| fuerte | 6 | 2 | 1.10E-05 | malo | 4 | 2 | 3.00E-05 |
| grande | 6 | 2 | 1.00E-04 | pobre | 5 | 2 | 5.00E-05 |
| largo | 5 | 2 | 2.22E-04 | puro | 4 | 2 | 2.50E-05 |
| lento | 5 | 2 | 1.70E-05 | rico | 4 | 2 | 2.32E-05 |
| rojo | 4 | 2 | 3.77E-05 | sabio | 5 | 2 | 1.38E-05 |
| seco | 4 | 2 | 1.80E-05 | serio | 5 | 2 | 3.96E-05 |
| suave | 5 | 1 | 3.00E-05 | tonto | 5 | 2 | 7.94E-06 |
| sucio | 5 | 2 | 6.78E-06 | triste | 5 | 2 | 3.36E-05 |
| verde | 5 | 2 | 3.78E-05 | vago | 4 | 2 | 5.00E-06 |

| **Noun Animal** | **Length** | **Syll.** | **Ngram Freq** | **Noun food** | **Length** | **Syll.** | **Ngram Freq** |
| --- | --- | --- | --- | --- | --- | --- | --- |
| 乳牛-cow | 2 | 2 | 1.00E-07 | 咖啡-coffee | 2 | 2 | 4.60E-05 |
| 企鵝-penguin | 2 | 2 | 1.40E-06 | 大蒜-garlic | 2 | 2 | 8.00E-07 |
| 兔子-rabbit | 2 | 2 | 3.30E-06 | 奶油-butter | 2 | 2 | 7.60E-07 |
| 海豹-seal | 2 | 2 | 5.10E-07 | 桃子-peach | 2 | 2 | 4.60E-07 |
| 熊-bear | 1 | 1 | 4.00E-06 | 水-water | 1 | 1 | 2.00E-04 |
| 狐狸-fox | 2 | 2 | 1.60E-06 | 沙拉-salad | 2 | 2 | 3.80E-06 |
| 狗-dog | 1 | 1 | 2.20E-05 | 洋蔥-onion | 2 | 2 | 1.50E-06 |
| 狼-wolf | 1 | 1 | 1.00E-05 | 牛奶-milk | 1 | 1 | 7.80E-06 |
| 猴子-monkey | 2 | 2 | 9.00E-06 | 玉米-corn | 1 | 1 | 1.40E-05 |
| 獅子-lion | 2 | 2 | 6.20E-06 | 米-rice | 1 | 1 | 6.00E-05 |
| 綿羊-sheep | 2 | 2 | 1.80E-06 | 糖-sugar | 1 | 1 | 8.40E-06 |
| 老虎-tiger | 2 | 2 | 7.00E-06 | 茶-tea | 1 | 1 | 2.20E-05 |
| 老鼠-mouse | 2 | 2 | 8.60E-06 | 葡萄-grape | 2 | 2 | 3.70E-06 |
| 蜘蛛-spider | 2 | 2 | 5.00E-06 | 蘋果-apple | 2 | 2 | 2.30E-05 |
| 螞蟻-ant | 2 | 2 | 4.60E-06 | 蛋-eggs | 1 | 1 | 7.70E-06 |
| 貓-cat | 1 | 1 | 9.10E-06 | 豆子-bean | 2 | 2 | 4.70E-07 |
| 馬-horse | 1 | 1 | 8.20E-05 | 辣椒-chili | 2 | 2 | 9.00E-07 |
| 鯨魚-whale | 2 | 2 | 1.80E-06 | 鳳梨-pineapp | 2 | 2 | 4.50E-07 |
| 鳥-bird | 1 | 1 | 1.30E-05 | 鹽-salt | 1 | 1 | 1.20E-05 |
| 龍-dragon | 1 | 1 | 2.00E-05 | 麵包-bread | 2 | 2 | 2.50E-05 |
| **Adj concrete** | **Length** | **Syll.** | **Ngram Freq** | **Adj abstract** | **Length** | **Syll.** | **Ngram Freq** |
| 乾的-dry | 2 | 2 | 1.00E-04 | 假的-fake | 2 | 2 | 4.00E-05 |
| 低的-low | 2 | 2 | 2.00E-04 | 傻的-silly | 2 | 2 | 4.00E-06 |
| 冷的-cold | 2 | 2 | 1.50E-05 | 壞的-bad | 2 | 2 | 4.30E-05 |
| 大的-large | 2 | 2 | 1.40E-03 | 好的-good | 2 | 2 | 6.00E-04 |
| 小的-small | 2 | 2 | 4.50E-04 | 帥的-handsor | 2 | 2 | 2.50E-06 |
| 快的-fast | 2 | 2 | 9.60E-05 | 忙的-busy | 2 | 2 | 1.50E-05 |
| 慢的-slow | 2 | 2 | 3.00E-05 | 怒的-angry | 2 | 2 | 5.50E-06 |
| 濕的-wet | 2 | 2 | 8.10E-06 | 懶的-lazy | 2 | 2 | 3.80E-06 |
| 熱的-hot | 2 | 2 | 5.50E-05 | 新的-new | 2 | 2 | 1.40E-03 |
| 甜的-sweet | 2 | 2 | 5.00E-06 | 病的-sick | 2 | 2 | 4.30E-04 |
| 瘦的-thin | 2 | 2 | 4.00E-06 | 瘋的-mad | 2 | 2 | 4.50E-06 |
| 短的-short | 2 | 2 | 5.00E-05 | 真的-real | 2 | 2 | 1.40E-04 |
| 硬的-hard | 2 | 2 | 2.00E-05 | 窮的-poor | 2 | 2 | 1.80E-05 |
| 胖的-fat | 2 | 2 | 6.30E-06 | 笨的-stupid | 2 | 2 | 2.20E-06 |
| 軟的-soft | 2 | 2 | 2.60E-05 | 累的-tired | 2 | 2 | 1.70E-05 |
| 輕的-light | 2 | 2 | 3.00E-05 | 美的-beautifu | 2 | 2 | 2.00E-04 |
| 酸的-sour | 2 | 2 | 3.80E-06 | 舊的-old | 2 | 2 | 8.00E-05 |
| 重的-heavy | 2 | 2 | 2.20E-04 | 貴的-expensi | 2 | 2 | 2.70E-05 |
| 長的-long | 2 | 2 | 3.00E-04 | 醜的-ugly | 2 | 2 | 6.30E-06 |
| 高的-tall | 2 | 2 | 5.00E-04 | 難的-difficult | 2 | 2 | 1.40E-04 |

**Table S4.**
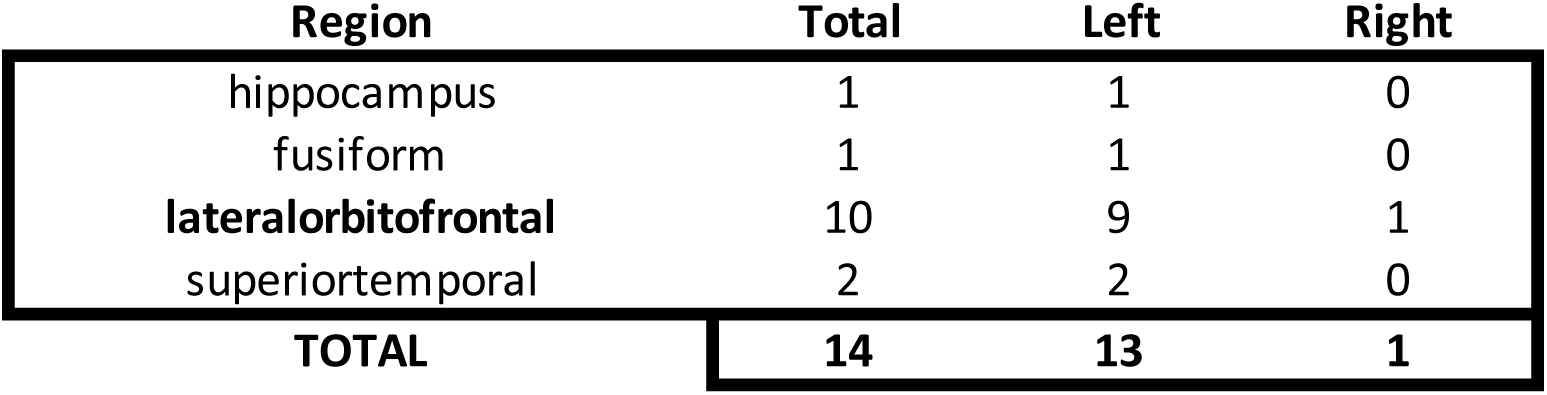
Distribution of electrodes that showed modulation by part of speech across brain regions. Significant regions showing lateralization shown in bold. (p<10^-5^, permutation test, n=10^6^ iterations, regions with less than 4 electrodes were excluded).

| Region | Total | Left | Right |
| --- | --- | --- | --- |
| hippocampus | 1 | 1 | 0 |
| fusiform | 1 | 1 | 0 |
| **lateralorbitofrontal** | 10 | 9 | 1 |
| superiortemporal | 2 | 2 | 0 |
| **TOTAL** | **14** | **13** | **1** |

**Table S5.** Distribution of nouns- versus adjective-preferring electrodes and electrodes that generalize for parts-of-speech versus those that do not. Distribution across different subjects of electrodes that are more noun enhanced (column 3) versus more adjective enhanced (column 4), and that of electrodes that generalize to nouns and adjectives (column 5) versus those that showed differences between noun subcategories or adjective subcategories (column 6).

| subject# | #Total | NounEnhanced<br>($\beta > 0$) | AdjEnhanced<br>($\beta < 0$) | Generalize | SubCategory<br>(=Total-Generalized) |
| --- | --- | --- | --- | --- | --- |
| 5 | 2 | 2 | 0 | 0 | 2 |
| 14 | 3 | 2 | 1 | 1 | 2 |
| 16 | 3 | 3 | 0 | 3 | 0 |
| 18 | 2 | 2 | 0 | 2 | 0 |
| 20 | 4 | 2 | 2 | 2 | 2 |
| **TOTAL** | **14** | **11** | **3** | **8** | **6** |

**Table S6.**
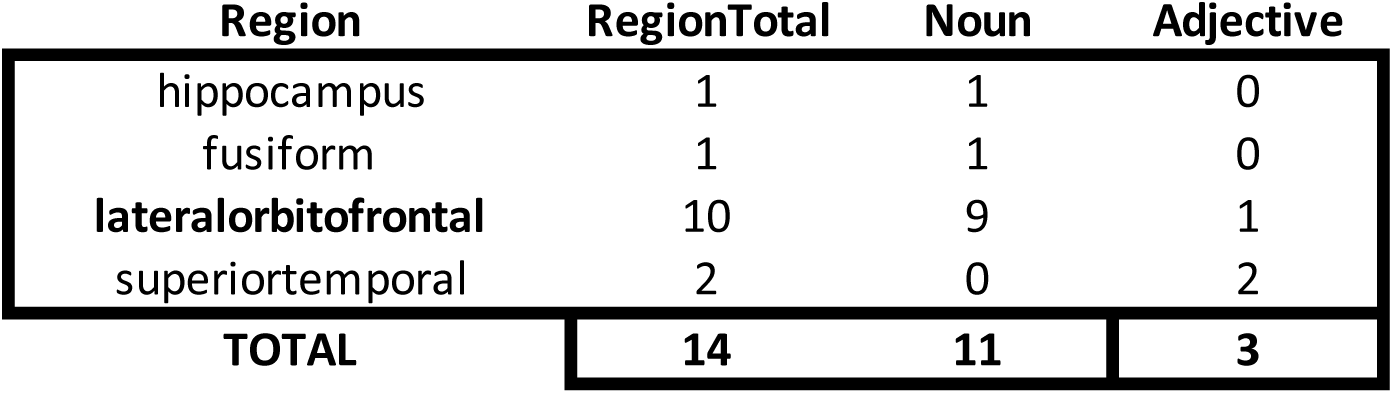
Distribution of nouns- versus adjective-preferring electrodes across brain regions. A permutation test combining all brain regions for these electrodes showed that LOF was significantly noun preferring. (p<10^-5^, permutation test, n=10^6^ iterations, regions with less than 4 electrodes were excluded).

**Table S7.**
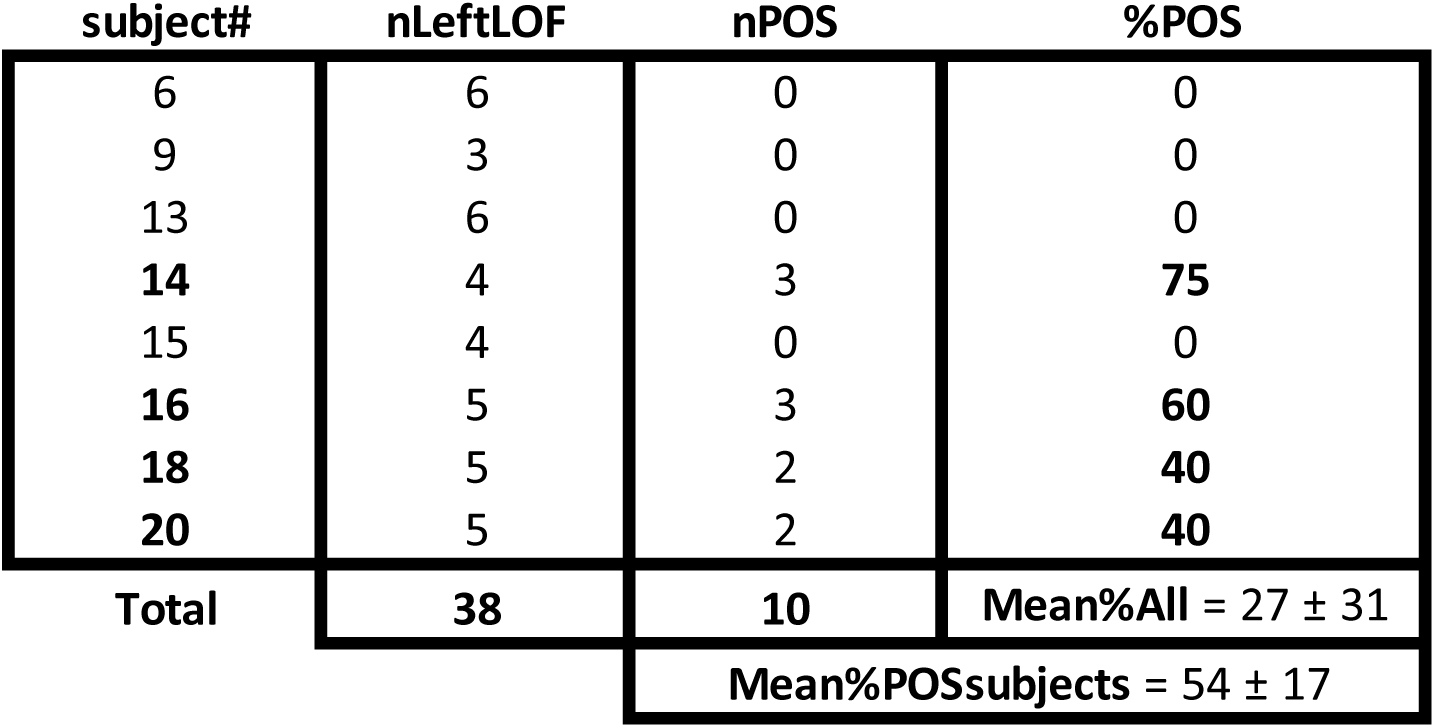
Distribution of part of speech encoding electrodes in the left lateralorbitofrontal cortex across subjects.

**Table S8.**
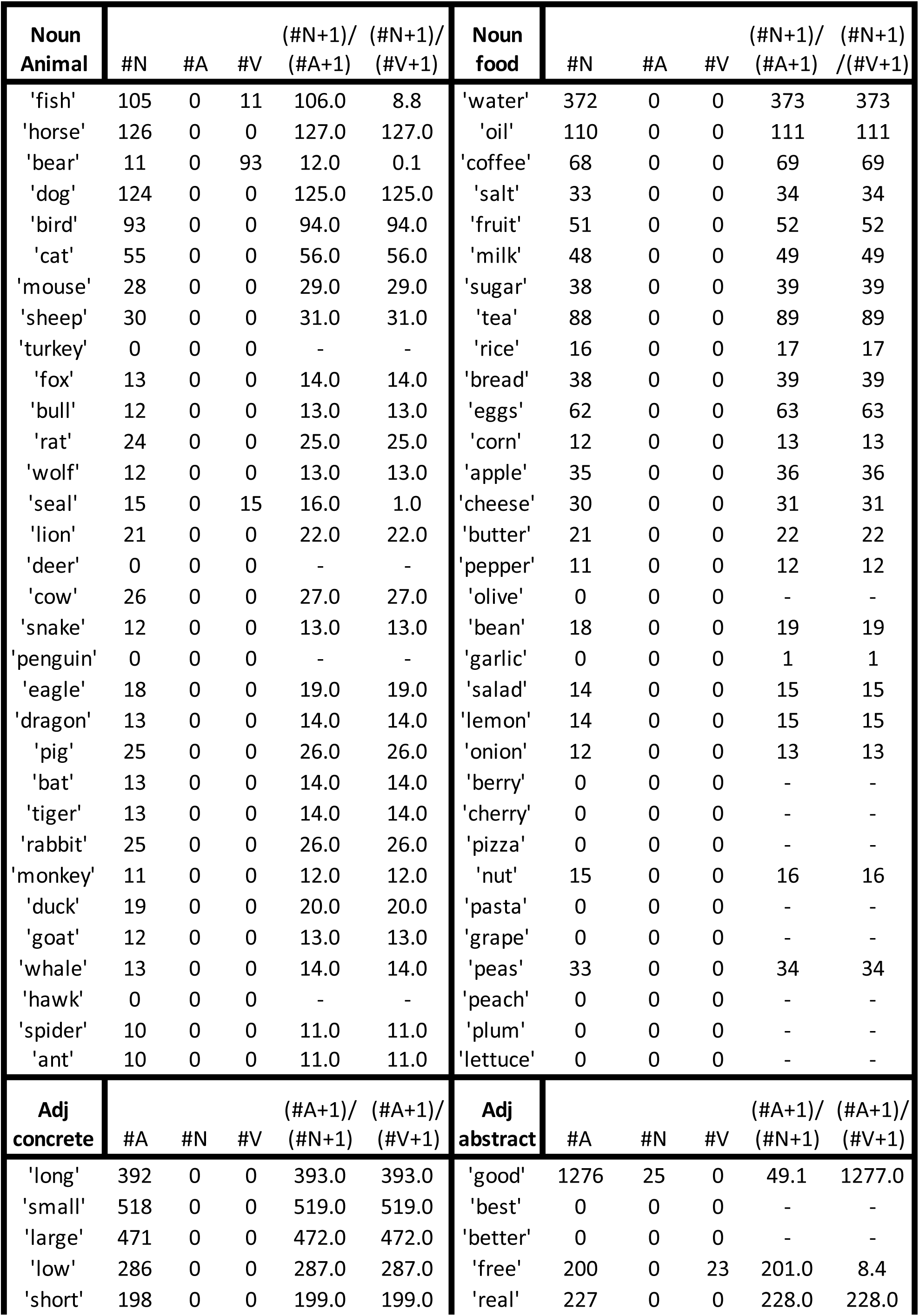
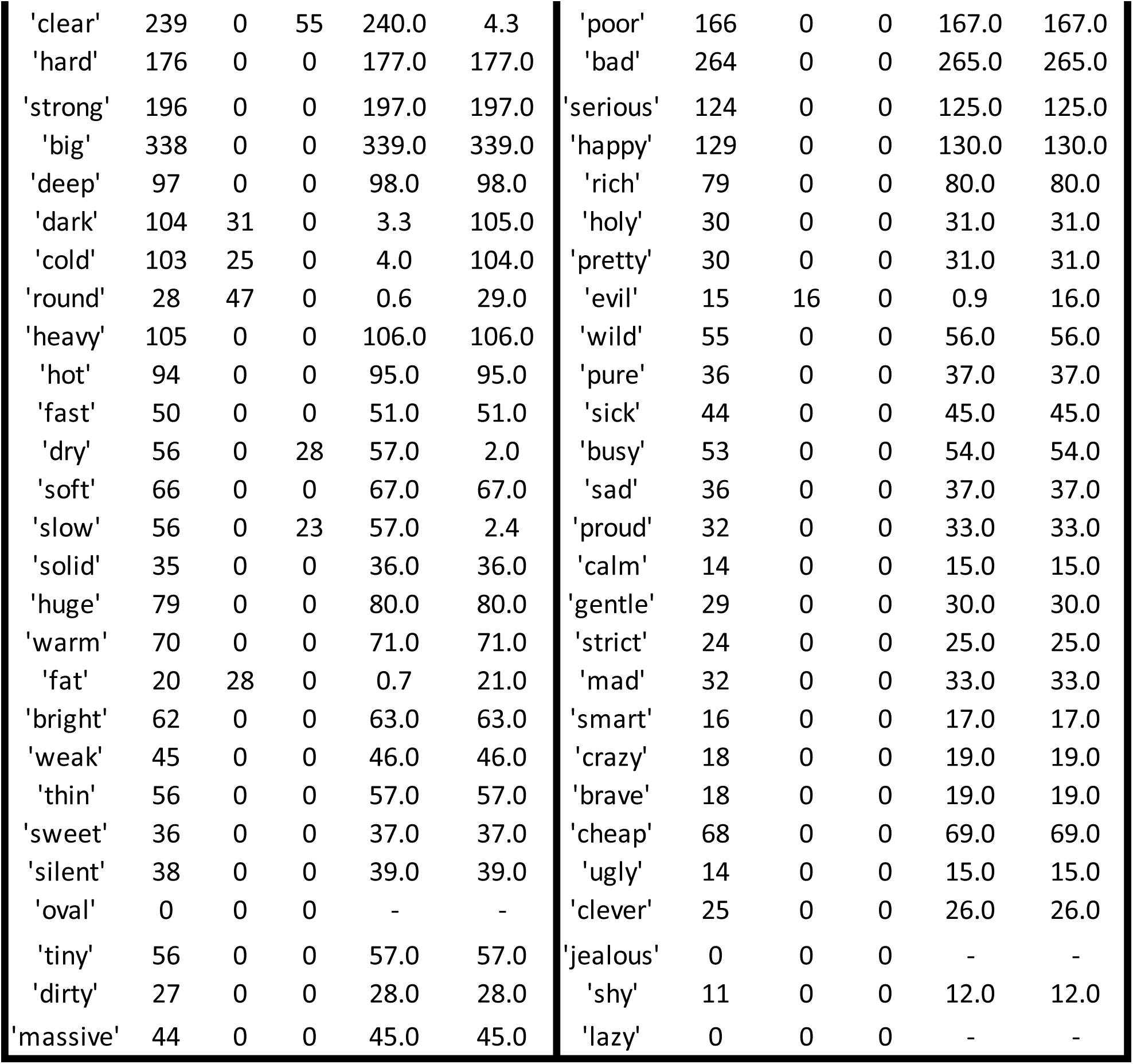
List of all the words used in the experiment, the number of times they occurred in the British National Corpus as a noun (#N), as an adjective (#A), or a verb (#V), and the ratios of their frequency of occurence in their assigned part of speech versus their usage in other parts of speech. Dashes indicate words that were missing in the corpus.

